# An age-preserving neuronal iSpheroid platform for modeling human aging and Alzheimer’s disease

**DOI:** 10.64898/2026.09.30.755853

**Authors:** Samhan Alsolami, Meiyan Wang, Thais Sarraf Sabedot, Upendra Singh, Brian Kirk, Lisa Mitchell, Melissa Hernandez, Nicholas Skvir, Chynna Bowman, Kotaro Oiwa, Ravi Agarwal, Marjan Mahmoudi, Michael S. Cuoco, Elise MacDonald, Samuel Cranston, Raffaella Lucciola, Daniela Boassa, Joseph R. Herdy, Jeffrey R. Jones, Kazuori Yoshikiyo, Longevity Consortium, Jerome Mertens, Julio Martinez-Trujillo, Lukasz Jaremko, George J. Murphy, Fred. H. Gage

## Abstract

Aging is the strongest risk factor for Alzheimer’s disease (AD), yet existing human neuronal models either erase age-associated features during reprogramming or fail to recapitulate three-dimensional organization of human neuronal tissue. Here, we establish directly induced neuronal spheroids (iSpheroids), a scalable three-dimensional platform generated by direct conversion of primary fibroblasts without pluripotency. iSpheroids generate long-lived, electrophysiologically active neurons from young, unimpaired aged, and sporadic AD donors while preserving donor-specific epigenetic age. Single-nucleus RNA sequencing revealed approximately 96% of cells acquired a neuronal identity and were distributed across multiple neuronal states, including a diversity of neurons. The platform supports incorporation of astrocytes and microglia and generation from non-human primate fibroblasts. AD iSpheroids spontaneously recapitulated neuronal degeneration, amyloid-β accumulation, progressive Tau pathology, and metabolic dysfunction. Integrated proteomic, metabolomic, and lipidomic profiling distinguished physiological aging from AD and revealed disease-associated metabolic remodeling. Oxaliplatin shifted AD iSpheroids toward oxidative metabolism, whereas a multi-compound screen identified trametinib as a modulator of inflammatory and age-associated programs.

**Graphical abstract:** 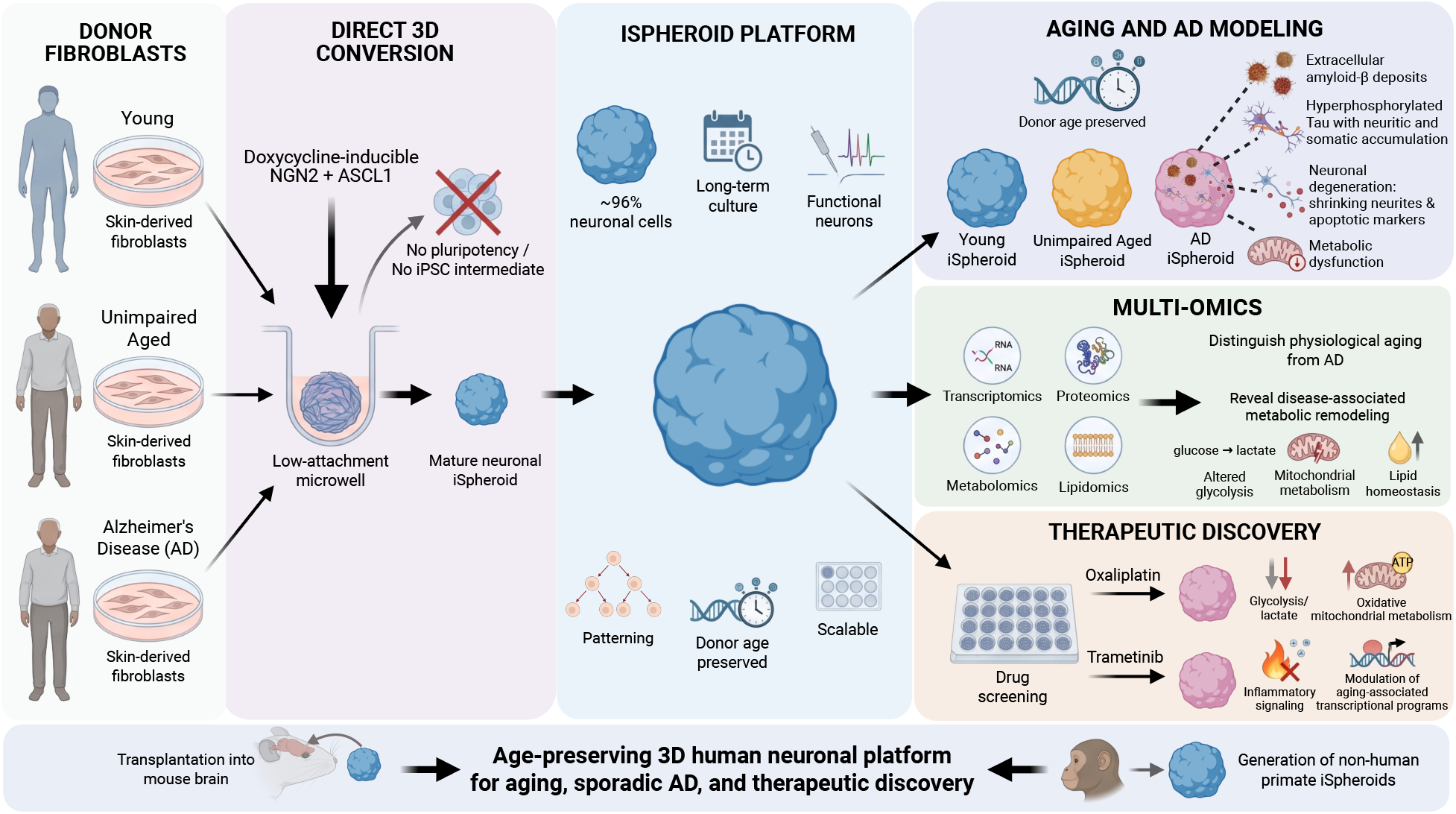

**Highlights:**

- Direct conversion generates age-preserving, functional human neuronal iSpheroids.
- iSpheroids recapitulate molecular and pathological hallmarks of AD.
- Multi-omics distinguishes physiological neuronal aging from AD.
- Pharmacological screening identifies reversible metabolic and inflammatory phenotypes.

## Introduction

Human aging is the greatest risk factor for most neurodegenerative disorders, including the sporadic form of Alzheimer’s disease (AD), yet the mechanisms by which aging renders human neurons susceptible to degeneration remain poorly understood^1–7^. Progressive deterioration of neuronal homeostasis during aging is accompanied by mitochondrial dysfunction^1,8–11^, impaired proteostasis^8,12,13^, oxidative stress^5,9,14^, chronic inflammation^15,16^, and declining synaptic plasticity^17,18^, all of which contribute to neuronal vulnerability and cognitive decline^5,19^. In AD, these age-associated changes are further exacerbated by amyloid-β accumulation^20–23^, tau pathology that includes hyperphosphorylation and mis-localization of Tau^24–26^, synaptic dysfunction^27,28^, and widespread neuronal degeneration^29–31^, ultimately leading to progressive impairment of memory and cognition. Although significant progress has been made in defining the molecular pathology of AD, understanding the molecular basis of how normal neuronal aging transitions into neurodegeneration - sometimes termed unsuccessful aging - remains a major challenge.

Human pluripotent stem cells have transformed the study of neurological diseases by providing *in vitro* renewable sources of patient-derived neurons. In particular, induced pluripotent stem cell (iPSC)-derived organoids have enabled modeling of human brain development^32–35^, neuronal circuit formation^36–38^, and disease-associated cellular interactions within three-dimensional (3D) environments that more closely resemble native brain tissue than conventional two-dimensional (2D) cultures^39–41^. These systems have become highly valuable for investigating neurodevelopmental disorders^35,42^ and have provided important insights into AD^43,44^, Parkinson’s disease (PD)^45,46^, and other neurological conditions^34^. Despite these advances, existing human *in vitro* models have important limitations for studying age-associated neurodegeneration; generating iPSC-derived brain organoids requires extensive cellular reprogramming from donor-specific fibroblasts, including subcloning and quality control, followed by prolonged differentiation, a process that can take several months before mature neuronal phenotypes emerge^36,47–49^. More importantly, iPSC-mediated reprogramming resets multiple hallmarks of cellular aging to an embryonic state, including epigenetic age, transcriptional programs, mitochondrial function, and other age-associated molecular features, thereby limiting the ability of iPSC-derived brain organoids to recapitulate age-associated neurodegenerative changes^47,50–53^. Consequently, many pathological phenotypes associated with aging must be artificially induced, reducing the ability to investigate the natural dynamics of disease mechanisms.

Direct neuronal conversion offers an attractive alternative by bypassing pluripotency and generating neurons directly from somatic cells while preserving important donor-specific aging signatures^50,54–63^. Through forced overexpression of neuronal factors such as NGN2 and ASCL1, directly converted induced neurons retain age-associated epigenetic, transcriptomic, and functional characteristics that are largely erased during iPSC reprogramming, making them particularly well suited for modeling sporadic neurodegenerative diseases in which aging is the principal risk factor. However, current direct conversion approaches are predominantly performed in 2D cultures for relatively short periods, often 21 days, limiting their ability to model neuronal aging and AD. In contrast, a 3D-based model can provide high neuronal density, neuronal connectivity, pathological progression, patterning toward mature neuronal states, scalability, reproducibility, and multicellular interactions that influence neuronal physiology and disease progression. Therefore, there is a need for a model that combines these advantages for modeling physiological and pathological neuronal aging.

Here, we developed a single-step direct neuronal conversion strategy that generates human iSpheroids, 3D structures composed predominantly of neuronal cells, from primary skin fibroblasts obtained from young, healthy aged, and sporadic AD donors. By bypassing pluripotency, fibroblasts rapidly self-assemble into highly reproducible 3D structures without centrifugation or embedding in extracellular matrices such as matrigel. These iSpheroids can be converted into long-lived, functionally active neurons while preserving donor-associated aging features. We show that iSpheroids recapitulate molecular, metabolic, and pathological features of human neuronal aging and AD and remain compatible with glial incorporation, *in vivo* transplantation, and pharmacological screening. Using this platform, we identify coordinated metabolic remodeling that distinguishes physiological aging from AD and show that disease-associated metabolic and transcriptional dysfunction can be partially reversed pharmacologically. Compared to conventional 2D systems, iSpheroids achieve high neuronal density without FACS purification and develop amyloid accumulation, neuronal degeneration, and progressive Tau pathology over time. Together, these findings establish iSpheroids as a scalable human neuronal platform for studying aging, modeling sporadic AD, and evaluating therapeutic interventions.

## Results

### A scalable, single-step fibroblast-derived iSpheroid platform generates mature human neurons from aged donors

To establish an iSpheroid model that preserves donor- and aging-specific biology, we developed a direct conversion strategy that bypasses the iPSC stage and generates 3D iSpheroids directly from primary human skin fibroblasts. These human fibroblasts were transduced with a lentiviral vector expressing doxycycline-inducible NGN2 and ASCL1 (see STAR methods) and allowed to aggregate by gravity into spheroid-like structures in U-bottom 96-well plates, followed by neuronal conversion for 21 days and maturation for more than 60 days (**Figure 1A**). This approach enabled the efficient generation of reproducible and uniformly sized neuronal spheroids from donors spanning young, healthy-aged, and AD donors. Consistent spheroid morphology was observed across independent donors, with reproducible diameters and highly uniform spheroid structures (**Figure 1B**; **Figure S1A**). iSpheroid self-assembly began with fibroblasts settling in the U-shaped wells at 0 h, followed by compaction within ∼5 h, with formation completed by 24 h as a cup-like structure termed an iSpheroid (**Figure S1B and Video S1**).

**Figure 1:**
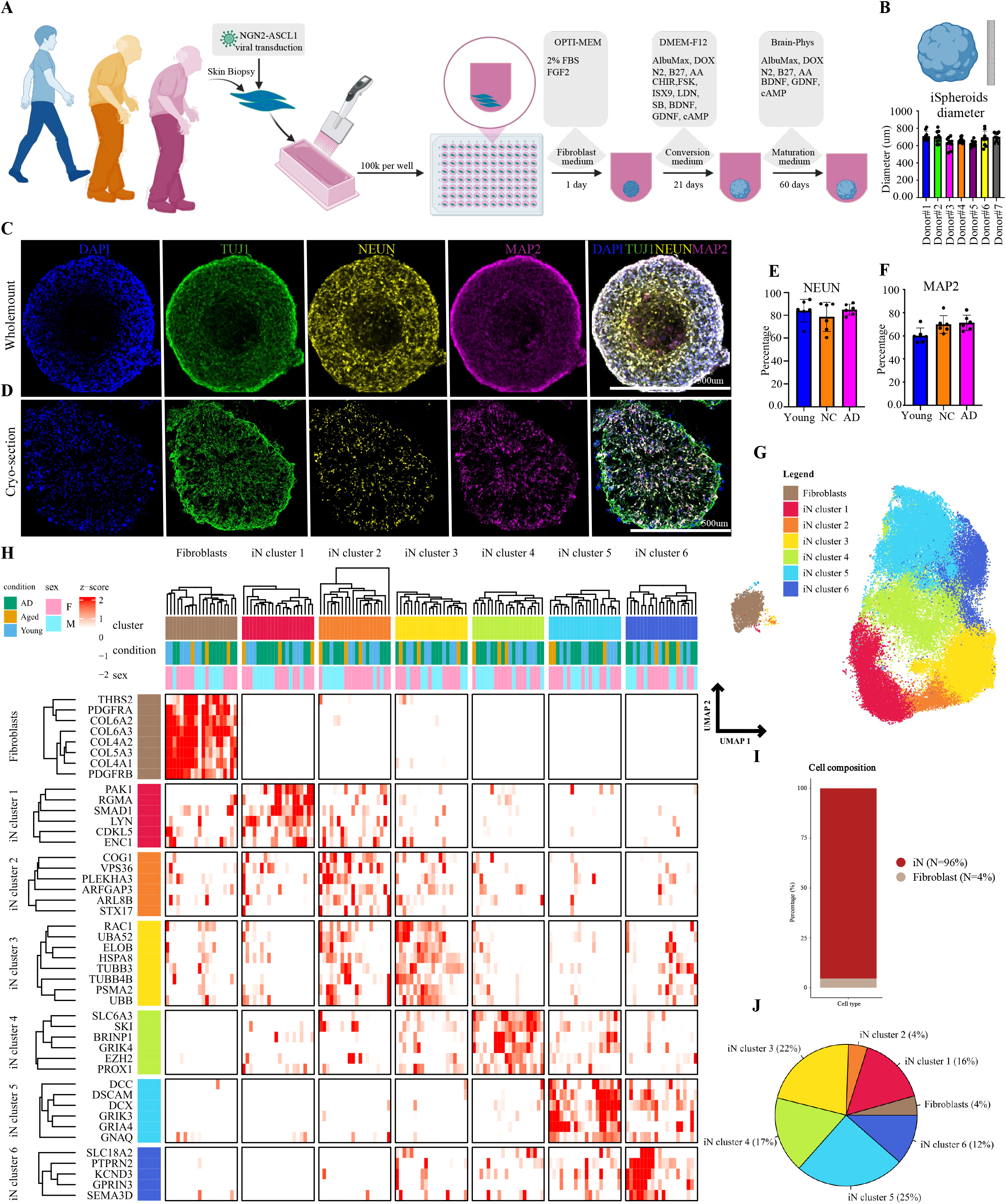
Generation, characterization, and single-nucleus transcriptomic profiling of human iSpheroids. **(A)** Schematic overview of iSpheroid generation from human skin biopsy-derived fibroblasts. Fibroblasts are transduced to generate stable cell lines carrying doxycycline-inducible neuronal transcription factors NGN2 and ASCL1. Cells are collected at a density of 10 million cells per 10 mL of medium and seeded into low-attachment 96-well plates at approximately 100,000 cells per well (100 μL per well). Fibroblasts are aggregated in fibroblast medium for ∼24 hours to form spheroids, followed by neuronal conversion in neuronal induction medium for ∼21 days and subsequent neuronal maturation for ≥60 days. **(B)** Diameter measurements of mature iSpheroids generated from seven independent donors (10 replicates). **(C-D)** Whole-mount and 20-μm cryosection immunofluorescence imaging of iSpheroids labeled with neuronal markers TUJ1, NEUN, and MAP2. Scale bar is 500 um. **(E-F)** Quantification of NEUN- and MAP2-positive cells across iSpheroids generated from young, healthy aged, and AD donors (n=6). **(G)** UMAP visualization of single-nucleus RNA sequencing data from iSpheroids derived from 20 donors spanning young, healthy aged, and AD donors groups, revealing multiple neuronal subpopulations and a small fibroblast subcluster (n=20). **(H)** Heatmap showing the top marker genes defining each cluster identified in the UMAP analysis. **(I)** Quantification of cell-type composition across all donors, demonstrating an average neuronal content of approximately 96% and a fibroblast content of approximately 4%. **(J)** A pie chart illustrating the proportional representation of cellular clusters within iSpheroids.

Histological assessment confirmed neuronal conversion within iSpheroids. Whole-mount imaging, cryosection analysis, and video imaging revealed widespread expression of the neuronal markers TUJ1, MAP2, and NEUN (**Figure 1C,D; Video S2**). Quantification further demonstrated that neuronal conversion efficiency remained robust across young, healthy aged, and AD donor-derived iSpheroids, with comparable proportions of MAP2- and NEUN-positive cells between donor groups (**Figure 1E,F**). Similar results were obtained using TUJ1 quantification (**Figure S1C**), demonstrating that advanced donor age and neurodegenerative status did not substantially impair direct neuronal conversion efficiency. Unlike MAP2 and TUJ1, NEUN relative to DAPI provided a more direct measure of neuronal abundance, with iSpheroids containing, on average, over 80% NEUN-positive cells (**Figure 1E).** Furthermore, long-term cultures maintained for up to 270 days retained neuronal marker expression and structural integrity in both healthy aged and AD-derived iSpheroids (**Figure S1D**), demonstrating that this iSpheroid platform supports prolonged neuronal maintenance suitable for modeling the neurobiology of aging and AD.

To define the cellular composition generated by direct conversion in iSpheroids, we performed single-nucleus RNA sequencing (snRNA-seq) on iSpheroids derived from 20 independent donors encompassing young, healthy-aged, and AD individuals. Unsupervised UMAP clustering identified six transcriptionally distinct neuronal populations together with a small fibroblast-like population composed of untransformed cells (**Figure 1G**). Cluster-defining marker genes confirmed robust molecular separation between neuronal states and residual fibroblasts (**Figure 1H**), whereas quantitative analysis showed that, on average, approximately 96% of cells acquired a neuronal identity, only ∼4% remained fibroblasts-like across all donors (**Figure 1I**). A pie chart summarizing cluster percentage composition further illustrates the diversity of neuronal sub-populations generated in this iSpheroid platform while emphasizing their predominant neuronal identity (**Figure 1J**). These findings were independently supported by donor-level quantification showing highly consistent neuronal enrichment across all 20 donors analyzed (**Figure S1E**). Gene Ontology (GO) enrichment further demonstrated that most of the neuronal clusters contained neuronal terms such as vesicle organization in cluster 2, axon guidance in cluster 3, and many synaptic terms in cluster 4,5, and 6 (**Figure S1F–L**). Together, unbiased snRNAseq assessment revealed that iSpheroids exhibited remarkable neuronal purity and consistency across multiple donors and across young, unimpaired, and AD groups.

To further resolve neuronal heterogeneity, we examined marker genes defining each transcriptional cluster. UMAP analysis identified six molecularly distinct neuronal subpopulations in addition to the residual fibroblast-like cluster (**Figure S2A**). Fibroblasts selectively expressed extracellular matrix-associated genes including ADAMTS6, GLIS3, and SPATS2L (**Figure S2B**), whereas each neuronal cluster exhibited a unique transcriptional signature characterized by genes such as ADAM19, NEURL2, ITPKB, CCDC39, ENTPD1, ZCCHC24, PLCG2, EGFL7, MT-ND4L, ELOVL2, CNR1, DAZL, NRCAM, SYNDIG1, KCTD16, WNT4, CACNA1D, and GPRIN3 (**Figure S2C–H**). These results indicate that direct neuronal conversion does not generate a single homogeneous neuronal population but instead produces multiple neuronal states with distinct molecular identities, consistent with progressive neuronal maturation and subtype diversification.

Because hippocampal vulnerability is a prominent feature of aging and AD^64–68^, we next asked whether iSpheroids generate neurons with hippocampus-like features. One neuronal cluster (cluster 4) was enriched for canonical hippocampal genes, consistent with a hippocampal transcriptional identity (**Figure S3A**). Immunofluorescence confirmed the presence of Prox1-positive neurons within mature iSpheroids; Prox1 is a well-established marker of dentate gyrus identity^69,70^ (**Figure S3B**). Further subdivision of cluster 4 resolved three transcriptionally distinct subpopulations that showed trends toward differential expression of PROX1 and GRIK4, although these differences did not reach statistical significance (**Figure S3C-E**), consistent with the presence of multiple hippocampal-like neuronal states. GRIK4 is an established marker of hippocampal CA3 identity^71^. These findings suggest that 3D direct conversion can generate specialized aged neuronal states relevant to brain regions that are particularly vulnerable during aging and neurodegeneration.

To determine whether neuronal conversion in iSpheroids influences cellular composition, we compared conventional 2D induced neurons with iSpheroids using snRNA-seq. Both systems produced a predominantly neuronal fate; however, iSpheroids exhibited a 6.25-fold reduction in fibroblast content compared to 2D culture, with neuronal-like cells comprising 96% of iSpheroids compared with 75% in 2D cultures (**Figure 2A–E**). Across donors, fibroblast content was significantly higher in 2D cultures than in iSpheroids, whereas neuronal content was significantly higher in iSpheroids than in 2D cultures (**Figure 2F-G**; p < 0.01 and p < 0.01, respectively). These findings indicate that iSpheroids preserve efficient neuronal conversion while potentially providing an environment that supports patterning toward specialized neuronal identities.

**Figure 2:**
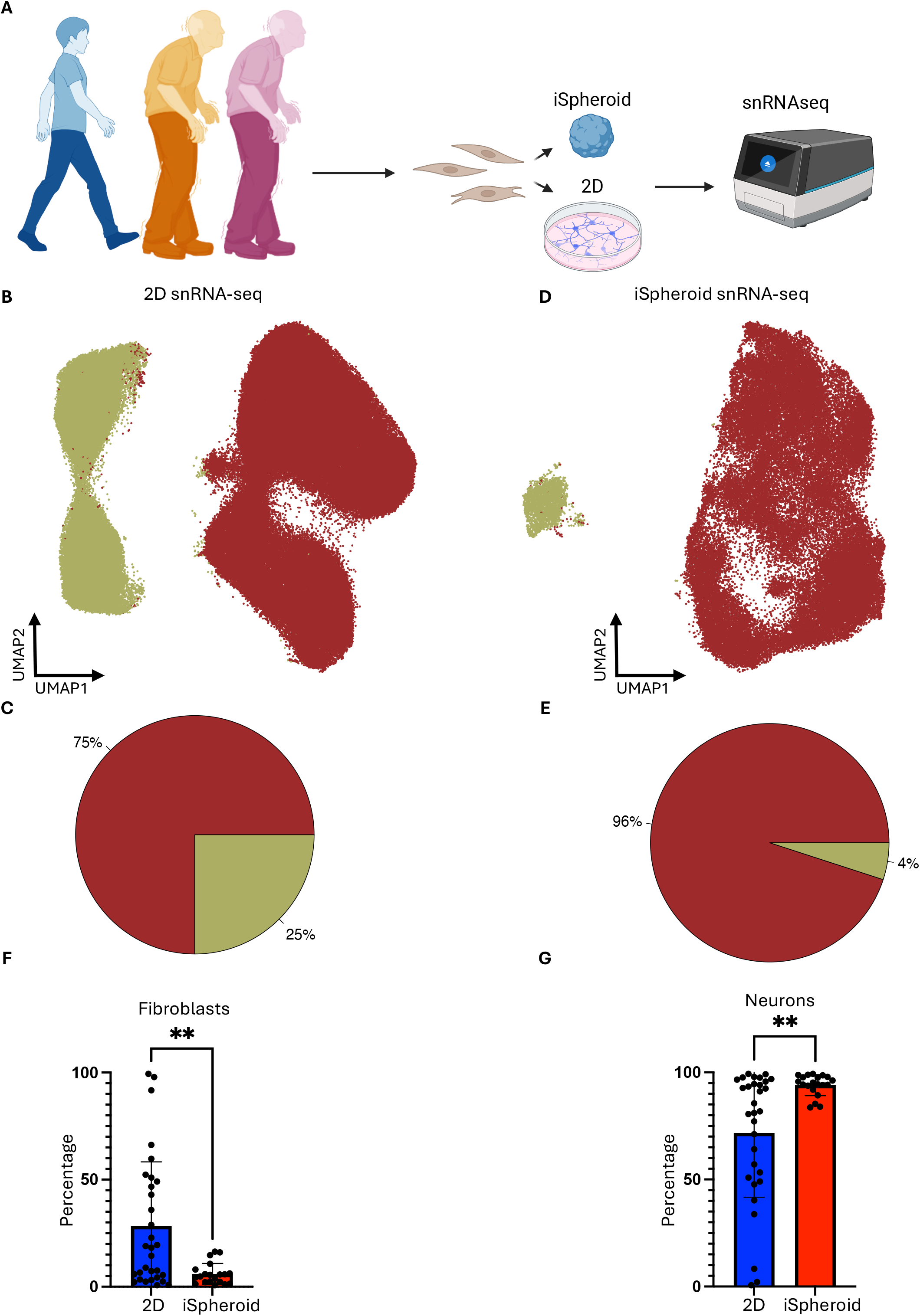
Benchmarking iSpheroids to 2D conventional neuronal conversion system. **(A)** Schematic showing the neuronal conversion of iSpheroid and 2D culture system followed by snRNAseq. **(B-C)** UMAP and pie chart showing average result of 2D-converted iNs (n=31). **(D-E)** UMAP and pie chart showing average result of iSpheroid-converted iNs (n=20). **(F-G)** Fibroblasts and neurons content comparing 2D and iSpheroid culture system. **, p < 0.01.

We next examined whether iSpheroids survive and maintain neuronal identity following transplantation into the adult mouse brain. Human iSpheroids were stereotaxically transplanted into mouse brain after in vitro neuronal conversion (**Figure S4A**). Histological analysis demonstrated extensive host-derived vascular infiltration throughout transplanted grafts (**Figure S4B, C**), indicating successful integration with the recipient microenvironment. Human cells remained readily identifiable by human nuclei staining and continued to express the mature neuronal markers MAP2 and NEUN following transplantation (**Figure S4D, E**), demonstrating long-term survival and maintenance of neuronal identity in vivo.

To evaluate the broader applicability of the platform, we generated iSpheroids from non-human primate (NHP), the common marmoset (Callithrix jacchus), using the same direct iSpheroid conversion strategy. Marmosets have emerged as an important nonhuman primate model of human aging^72^. Marmoset fibroblasts efficiently formed spheroids with uniform morphology and exhibited robust TUJ1 expression throughout the iSpheroids (**Figure S5A–C**), demonstrating that the approach is not restricted to human cells but is readily transferable to other NHP species.

A critical requirement for disease modeling is generating functionally mature neurons. To assess neuronal physiology, we performed whole-cell patch-clamp recordings together with multielectrode array (MEA) analyses (**Figure S6A**). iSpheroid-derived neurons generated repetitive action potentials under current-clamp conditions and exhibited spontaneous synaptic activity during voltage-clamp recordings (**Figure S6B,C**). Quantitative electrophysiological analyses demonstrated mature intrinsic membrane properties across neurons derived from young, healthy aged, and AD donors (**Figure S6D–K**). All groups showed a wide range of resting membrane potentials (RMPs) likely due to different stages of maturation^73^. Importantly, no significant differences were observed among groups, except for a significantly greater spike afterhyperpolarization (AHP) in AD donors than in healthy aged donors (**Figure S6H;** p<0.05).

MEA recordings further revealed spontaneous population-level electrical activity (**Figure S6L-M; Video S3**), showing that neurons form functional synapses that produce network bursts^74^ (**Figure S6N**). Calcium imaging independently confirmed spontaneous neuronal activity (**Video S4**). Scanning electron microscopy revealed synaptic structures within iSpheroids (**Figure S6O**). Together, these findings demonstrate that iSpheroids form physiologically active neuronal networks.

Finally, we expanded the platform to incorporate donor-matched glial cells. Donor-specific astrocytes and erythromyeloid progenitor-derived microglia were derived from iPSCs and integrated during iSpheroid generation (**Figure S7A**). Immunostaining confirmed successful incorporation of S100B-positive astrocytes and IBA1-positive microglia (**Figure S7B-C**); snRNA-seq identified neuronal, astrocytic, oligodendrocyte progenitor (OPC-like cells), microglial, fibroblast, and intermediate cell populations within glia-integrated iSpheroids (**Figure S7D-E**). While the presence of OPC-like cells was unexpected, previous studies suggest that astrocytes can acquire OPC-like features under neuronal like conversion conditions^75^. Further analysis demonstrated transcriptionally distinct fibroblast-like, neuronal, glial-like, OPC-like, microglial progenitor, mature microglial, and undifferentiated populations defined by lineage-specific marker genes across nine donors (**Figure S8A-B**), illustrating the cellular complexity that can be achieved within the platform. Successful seeding of iSpheroids with glial cells can be influenced by the timing of seeding relative to the age of both the glial cells and iSpheroids, as well as by glial cell density.

Collectively, these results establish iSpheroids as a single step, scalable, reproducible, and versatile 3D human neuronal system that efficiently generates mature and functionally active neurons from aged individuals while remaining compatible with multicellular modeling of the aging and AD brain.

### iSpheroids preserve human aging while recapitulating molecular and pathological hallmarks of AD

A central limitation of pluripotent stem cell-based models of late-onset neurodegenerative disease is the loss of donor age during cellular reprogramming. We therefore asked whether direct conversion into iSpheroids could preserve donor-specific aging signatures while capturing molecular and pathological features associated with AD. To address this question, we generated iSpheroids from young, healthy aged/unimpaired (NC), and AD donors and characterized them using snRNA-seq, DNA methylation profiling, immunofluorescence, biochemical assays, and scanning electron microscopy (SEM) (**Figure 3A**).

**Figure 3:**
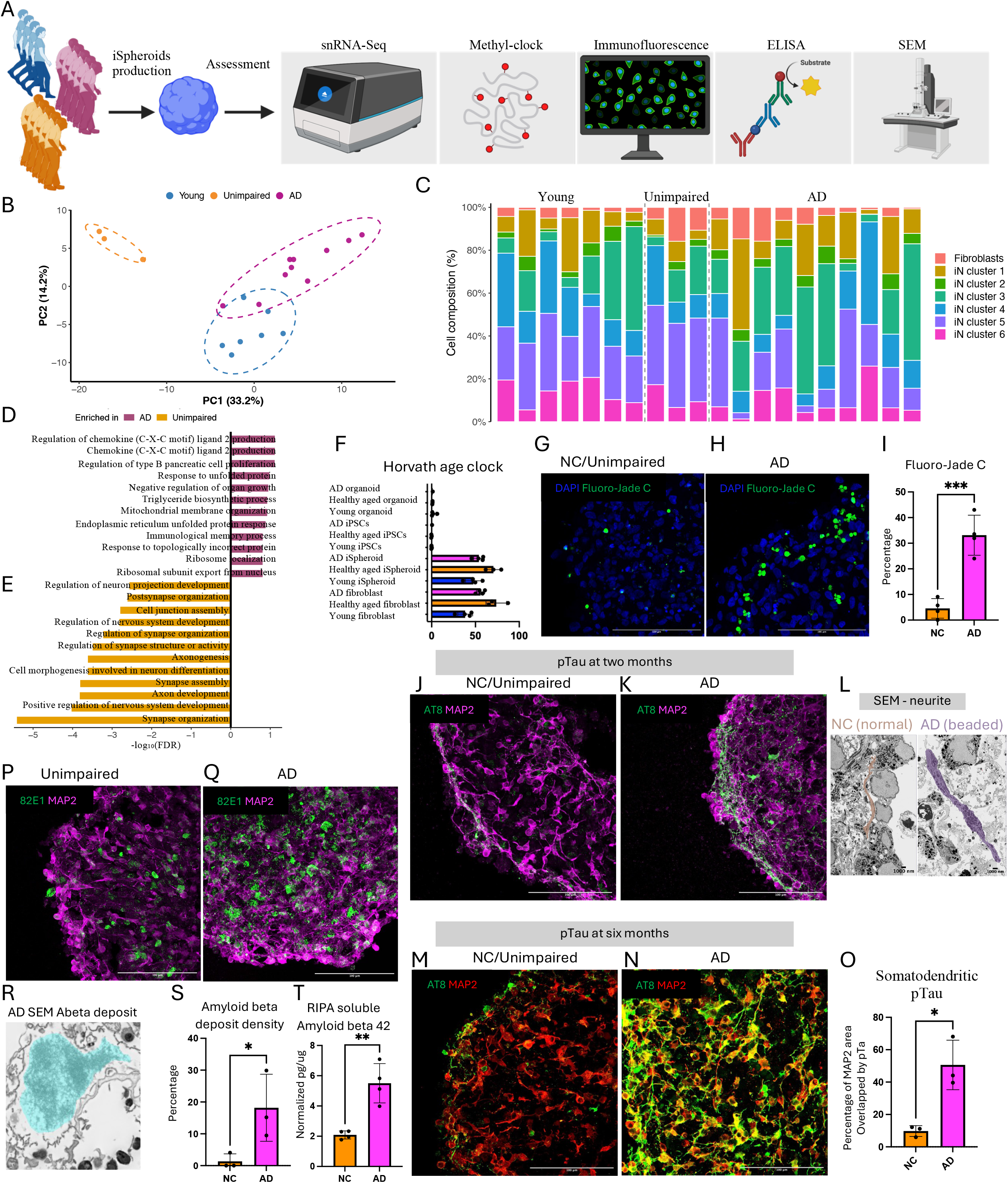
iSpheroids capture donor-specific neuronal aging and AD. **(A)** Schematic showing iSpheroid production from young, healthy aged/unimpaired (NC), and AD donors, followed by omics, immunofluorescence, ELISA, and scanning electron microscopy analyses. **(B)** Supervised PCA analysis of snRNA-seq data on 20 donors from young, unimpaired, AD donors, showing clear separation among donor groups. **(C)** Percentage composition of snRNA-seq clusters for each donor across young, unimpaired, and AD groups. (**D-E**) GO terms analysis comparing AD and unimpaired donors. (**F**) Methylation-clock analysis of donors representing 4 young, 3 unimpaired/healthy aged, and 4 AD donor groups at the fibroblast, iSpheroid, iPSC, and iPSC-derived organoid stages. The analysis shows that iSpheroids maintain donor-specific age, in contrast to iPSCs and iPSC-derived organoids. (**G–I**) Neurodegeneration assay using Fluoro-Jade C staining showing representative unimpaired (**G**) and AD (**H**) iSpheroids and quantification across four independent donors from each group (**I**). ***, p < 0.001. Scale bar is 100 um, (n=4). (**J–L**) Immunofluorescence assessment of Tau hyperphosphorylation in aged iSpheroids using the AT8 antibody, showing representative images of unimpaired (**J**) and AD (**K**) iSpheroids and scanning electron microscopy showing the bead-on-a-string morphology of dystrophic neurites found in AD alongside normal neurite from unimpaired donor (**L**). Scale bar is 100 um for immunofluorescent images, and 1000 nm for SEM. (**M–O**) AT8 staining of 6-month-old iSpheroids showing representative images of an unimpaired donor (**M**) and an AD donor (**N**) and quantification of somatic pTau localization by calculating the percentage of MAP2-positive area co-occupied by AT8 signal (**O**) on 3 donors per category. *, p < 0.05. Scale bar is 100 um. (**P–S**) Immunofluorescence staining of amyloid-β (Aβ) deposits using the 82E1 antibody (green) and MAP2 (magenta), showing representative images of an unimpaired donor (**P**) and an AD donor (**Q**), scanning electron microscopy of amyloid-β deposits (**R**), and quantification of the percentage of MAP2-positive area occupied by Aβ staining (**S**) on 3 donors per category. *, p < 0.05. Scale bar is 100 um for images, and 1000 nm for SEM. (**T**) ELISA-based measurement of Aβ42 in the RIPA-soluble fraction of iSpheroids from unimpaired and AD donors, 4 donors per category, normalized to the total protein content of the iSpheroids. **, p < 0.01.

We first examined whether donor age and disease status were reflected at the transcriptional level. snRNA-seq analysis revealed distinct distributions of cells derived from young, unimpaired, and AD donors (**Figure S9A**). Supervised PCA separated young, unimpaired, and AD donors (**Figure 3B**); the top 200 group-variable genes further distinguished these groups at the transcriptional level (**Figure S9B**). Analysis of cellular composition revealed a progressive depletion of mature neuronal populations in AD, particularly clusters 4, 5, and 6 (**Figure 3C and S9C;** *, p < 0.05), suggesting that AD pathology is accompanied by changes in the representation of mature neuronal states.

Pathway-level analyses further revealed molecular transitions associated with aging and AD. Kyoto Encyclopedia of Genes and Genomes (KEGG) enrichment analysis on mature neuronal clusters in snRNA-seq identified pathways distinguishing unimpaired aged donors from young donors (**Figure S10A**). It is well established that one of the strongest hallmarks of aging is loss of proteostasis^8,13,76^. Consistent with the literature, KEGG analysis revealed that young donors were enriched for terms such as proteolysis, metabolism and DNA repair relative to the unimpaired donors (**Figure S10A).** Comparison of AD with unimpaired donors identified broad disease-associated pathway alterations that include the enrichment of chemokine related terms, ER and mitochondria related terms, unfolded protein response, and lipid synthesis related terms in AD compared to unimpaired donors (**Figure 3D**), and the enrichment of rather positive terms in unimpaired donors compared to AD, which include synapse developments and neuronal maturation terms (**Figure 3E)**. In line with an enrichment of protein processing in ER and mitochondria terms (**Figure 3D**), SEM analysis showed dilated ER and abnormal mitochondria morphology in AD compared to the unimpaired (**Figure S10B**). Consistent with these pathway-level changes, few genes previously implicated in AD biology, including CDK5, PTPN1, and PDK1, were significantly increased in AD relative to unimpaired donors (**Figure S10C-E;** p < 0.01, p < 0.05, and p < 0.05, respectively). All of these genes have been implicated in AD biology: CDK5 contributes to Tau phosphorylation and amyloid-β generation^77–79^,, PTPN1 contributes to neuronal insulin resistance^80,81^, and PDK1 modulates neuronal responses to amyloid-β toxicity^82^. Moreover, AD iSpheroids displayed increased expression of apoptosis-related genes in mature neuronal clusters in snRNA-seq compared to unimpaired donors (**Figure S11A**), accompanied by enrichment of multiple apoptosis and cell death-associated gene sets (**Figure S11B**). Together, these findings indicate that iSpheroids capture molecular changes found in young, unimpaired and AD groups.

We next asked whether iSpheroid preserved the biological age of the donor. Epigenetic age was estimated with the Horvath pan-tissue clock, a 353-CpG elastic-net predictor trained to estimate chronological age across human tissues^83–85^. This DNA methylation clock analysis was performed across matched fibroblasts, directly converted iSpheroids, iPSCs, and iPSC-derived organoids. iSpheroids retained donor-specific epigenetic age signatures resembling those of the parental fibroblasts, whereas reprogramming to pluripotency markedly reduced these age-associated signatures in both iPSCs and their derived organoids (**Figure 3F**). Unexpectedly, methylome profiling revealed that AD neurons exhibited a young-like epigenetic state relative to age-matched unimpaired neurons, suggesting that AD-associated epigenetic remodeling diverges from, rather than simply extends, the normal neuronal aging trajectory (**Figure 3F**). Interestingly, previous epigenomic analyses of human brain tissues have similarly shown that AD does not simply extend the normal aging trajectory^86^. It has also been reported that H4K16ac changes during normal aging and AD were negatively correlated, with predominant H4K16ac gains during healthy aging but losses in AD, suggesting that AD disrupts rather than simply accelerates normal age-associated epigenetic remodeling^86^. Thus, unlike models that pass through a pluripotent intermediate, iSpheroids preserve an important molecular feature of human aging.

We next determined whether these age- and disease-associated molecular states were accompanied by neurodegenerative phenotypes. Consistent with the snRNA-seq findings (**Figure S11A-B),** Fluoro-Jade C staining, an established marker of neurodegeneration^87–92^, has revealed increased neuronal degeneration in AD iSpheroids compared with unimpaired controls across independent donors (**Figure 3G–I**; p < 0.001). AD iSpheroids also exhibited altered extracellular lactate levels (**Figure S11C;** p < 0.05), consistent with disease-associated metabolic remodeling and a Warburg-like phenotype previously described in AD^93–95^. Together with the enrichment of apoptotic transcriptional programs in AD, these findings link the molecular disease signatures detected by snRNA-seq to measurable cellular degeneration and metabolic dysfunction.

Tau pathology was also recapitulated in AD iSpheroids. AT8 immunostaining, an established approach for assessing Tau pathology^96–101^, revealed prominent hyperphosphorylated Tau in AD iSpheroids compared with unimpaired controls, frequently displaying a beads-on-a-string morphology reported in AD^62,102,103^ (**Figure 3J-K**). SEM independently revealed dystrophic neurites characterized by accumulation of organelles in axons (**Figure 3L**). After prolonged culture of iSpheroids for six months, AD iSpheroids exhibited pronounced redistribution of phosphorylated Tau toward somatodendritic compartments, as reflected by increased overlap between AT8- and MAP2-positive regions, consistent with the somatodendritic redistribution of phosphorylated Tau reported in AD postmortem tissues^104–107^ (**Figure 3M-O**; p < 0.05). Gel-based analysis also detected both 3R and 4R Tau isoforms across young, unimpaired, and AD iSpheroids and revealed group-associated differences in isoform abundance, with AD iSpheroids showing a trend toward a lower 4R Tau level (Figure S11D). The presence of 4R Tau is a well-established feature of adult neurons^108–110^. Together, these observations indicate that iSpheroids support the emergence and temporal progression of multiple Tau-associated phenotypes.

Additionally, we examined whether iSpheroids develop amyloid beta-like pathology. Amyloid pathology was assessed using 82E1, an Aβ N-terminal antibody preferentially recognizing the free Asp1 terminus of Aβ1-x with minimal cross-reactivity to APP^111,112^. Immunofluorescence using 82E1 antibody revealed extracellular Aβ-positive deposits in AD iSpheroids that were less prominent in unimpaired aged controls (**Figure 3P-Q**). SEM independently identified extracellular plaque-like structures with morphology consistent with the deposits detected by 82E1-based immunofluorescence (**Figure 3R**). Quantification of 82E1-positive signal relative to MAP2 demonstrated a significant increase in Aβ burden in AD iSpheroids (**Figure 3S**; p < 0.05). Consistent with the imaging results, ELISA of the RIPA-soluble fraction revealed increased soluble Aβ42 levels in AD relative to unimpaired iSpheroids after normalization to total protein content (**Figure 3T**; p < 0.01).

Collectively, these data demonstrate the ability of iSpheroids to preserve donor-specific molecular aging while simultaneously capturing transcriptional, metabolic, neurodegenerative, Tau, and amyloid-associated features of AD. In addition, AD donors displayed a loss of mature neuronal states, induction of AD-associated genes, and enrichment of pathways associated with AD and neurodegeneration. Thus, iSpheroids provide an age-preserving human neuronal model capable of resolving both aging- and disease-associated traits.

### Integrated multi-omics reveals metabolic remodeling that distinguishes physiological aging from AD

To further define the molecular programs that distinguish physiological neuronal aging from AD, we performed integrated multi-omics profiling of iSpheroids using lipidomics, metabolomics, and proteomics (**Figure 4A**). These complementary approaches enabled assessment of lipid composition, intracellular metabolism, and protein abundance across young, unimpaired, and AD donors.

**Figure 4:**
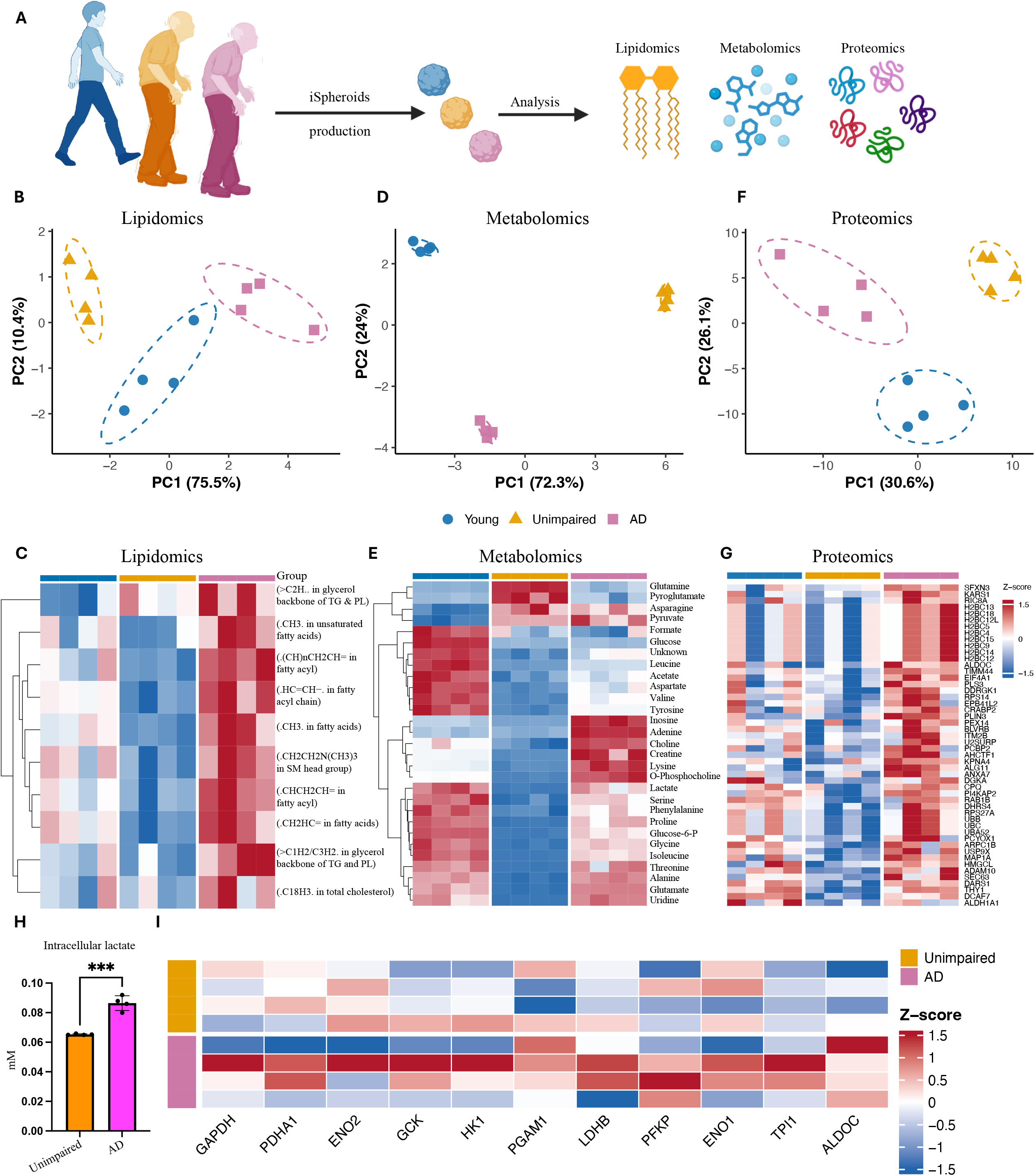
Multi-omics profiling of human iSpheroids across aging and AD. **(A)** Schematic overview of iSpheroid generation followed by comprehensive multi-omics analyses, including proteomics, metabolomics, and lipidomics, with four donors representing each category. **(B–C)** Principal component analysis (PCA) and heatmap of lipidomic datasets obtained from iSpheroids derived from young (blue), healthy aged (orange), and AD (magenta) donors, demonstrating donor group-specific molecular signatures (n=4). **(D–E)** Principal component analysis (PCA) and heatmap of metabolomic datasets across young, healthy aged (unimpaired), and AD donor-derived iSpheroids, highlighting molecular signatures associated with aging and neurodegeneration (n=4). **(F–G)** Principal component analysis (PCA) and heatmap of proteomic datasets across young, healthy aged (unimpaired), and AD donor-derived iSpheroids, highlighting molecular signatures associated with aging and neurodegeneration (n=4). **(H)** Metabolomics-based intracellular assessment of lactate accumulation in unimpaired and AD donors. ***, p < 0.001, (n=4). **(I)** Heatmap showing the relative abundance of glycolytic pathway proteins across unimpaired and AD donor-derived iSpheroids. Rows represent individual donors, grouped as unimpaired and AD, while columns represent glycolytic proteins. Color intensity indicates row-wise Z-score– normalized abundance, with red indicating higher abundance and blue indicating lower abundance.

We first examined lipidomic differences across donor groups. Principal component analysis and heatmap-based assessment separated young, unimpaired, and AD iSpheroids (**Figure 4B–C and Figure S12A–C**). AD iSpheroids showed increased abundance of multiple lipid species, whereas young donors displayed a similar but less pronounced pattern. In contrast, unimpaired aged donors showed comparatively lower lipid abundance across these measurements. These findings indicate that lipid composition differs between physiological aging and AD, with AD characterized by increased accumulation of intracellular lipids. This result is in line with reported findings that AD neurons accumulate lipids^113–115^, further demonstrating that iSpheroids capture pathological features of AD.

Metabolomic profiling similarly distinguished young, unimpaired, and AD iSpheroids (**Figure 4D–E and Figure S13A–F**). Young and AD donors showed higher abundance of multiple intracellular metabolites compared with unimpaired donors, paralleling the pattern observed by lipidomics. We further performed metabolomic analysis of conditioned media to determine whether these intracellular differences were accompanied by changes in extracellular metabolites. Although conditioned-media metabolomics did not clearly separate young, unimpaired, and AD groups, several metabolites showed group-associated patterns consistent with those observed intracellularly (**Figure S14A–B and Figure S15A–H**). Together, the lipidomic and metabolomic datasets identify a distinct metabolic state in unimpaired aged neurons compared with both young and AD iSpheroids.

We next examined whether these differences were also reflected at the protein level. Proteomic analysis separated young, unimpaired, and AD (**Figure 4F–G and S16A-D**). Differential protein abundance analysis further identified disease- and age-associated changes across these groups. Compared with unimpaired donors, AD iSpheroids showed increased abundance of several proteins implicated in AD and neurodegeneration, including ITM2B, a regulator of APP processing^116,117^ (**Figure 4G**). Further comparison of AD with unimpaired donors identified increased abundance of AD-associated proteins, including ADAM10, which participates in APP processing^118–120^ (**Figure S16C**). In contrast, comparison of unimpaired with young donors identified increased abundance of aging-associated proteins, including GPNMB, POSTN, and SPON1, in unimpaired iSpheroids (**Figure S16A**). GPNMB has been shown in many publications to be well correlated well with aging^121–126^, further supporting our iSpheroids model’s ability to capture human aging. Also, GPNMB is enriched in aging and disease-associated microglia and can undergo ectodomain shedding, generating soluble GPNMB that has emerged as a potential biomarker of aging, with microglia representing a major cellular source of GPNMB in the brain^127^.

Pathway-level analysis further distinguished the molecular states associated with aging and AD. Gene ontology (GO) analysis showed that young donors were enriched for terms related to cell cycle regulation, whereas unimpaired donors were enriched for terms associated with mitochondrial function (**Figure S16B**). In contrast, comparison of unimpaired and AD donors revealed enrichment of terms related to nucleosome assembly and amyloid fibril formation in AD, whereas unimpaired donors were enriched for terms associated with biosynthetic and metabolic processes (**Figure S16B**).

Because altered energy metabolism is associated with AD, we next examined glycolytic activity in iSpheroids. NMR-based measurement of intracellular lactate revealed increased lactate accumulation in AD donors compared with unimpaired donors (**Figure 4H**; p < 0.001). Consistent with this observation, proteomic analysis showed increased abundance of multiple glycolysis-associated enzymes in AD compared to unimpaired donors, including GAPDH, PDHA1, ENO2, GCK, HK1, PGAM1, LDHB, PFKP, ENO1, TPI1, and ALDOC (**Figure 4I**).

Together, these multi-omics analyses revealed molecular differences between physiological aging and AD across lipid, metabolite, and protein levels. Unimpaired aged iSpheroids displayed lower intracellular lipid and metabolite abundance, whereas AD iSpheroids showed increased lipid accumulation and altered metabolic profiles. Furthermore, in these multi-omics analyses, young donors often show intermediate features between unimpaired and AD donors. These findings indicate that iSpheroids capture metabolic and proteomic remodeling associated with both physiological neuronal aging and AD.

### Oxaliplatin reverses metabolic dysfunction in AD iSpheroids

To determine whether a disease-associated phenotype in iSpheroids could be pharmacologically perturbed, we targeted the metabolic dysfunction observed in AD cultures. Having established that AD iSpheroids capture extensive metabolic remodeling, including a Warburg-like phenotype (**Figure S12C and Figure 4H-I**), we asked whether this abnormal state could be shifted toward oxidative metabolism. AD iSpheroids from multiple donors were treated with the glycolysis-suppressing agent oxaliplatin for two days and then evaluated by extracellular lactate measurement, bulk RNA sequencing, and Seahorse analysis (**Figure 5A**). Clinically, oxaliplatin is used to treat colorectal cancer, and its therapeutic response has also been linked to suppression of glycolytic metabolism^128^.This integrated design enabled parallel assessment of metabolic output, transcriptional remodeling, and mitochondrial function following treatment.

**Figure 5:**
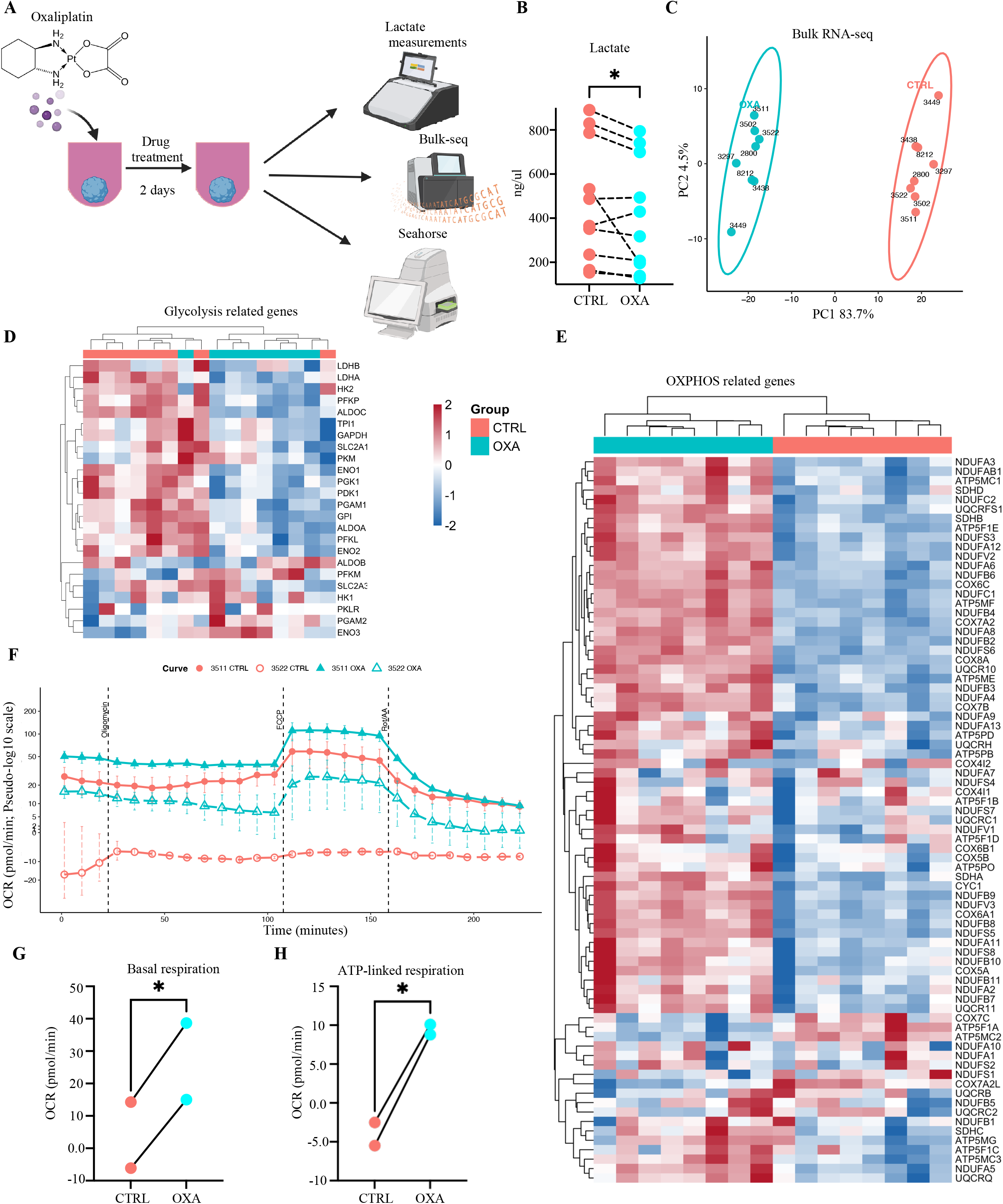
Oxaliplatin reverses metabolic dysfunction in AD iSpheroids. **(A)** Schematic overview of the experimental design in which AD donor-derived iSpheroids were treated with Oxaliplatin for two days, followed by extracellular lactate measurements, bulk RNA sequencing analysis, and seahorse analysis. **(B)** Quantification of lactate metabolism in iSpheroids derived from 10 AD donors demonstrating that Oxaliplatin treatment suppresses the Warburg-like metabolic phenotype, consistent with reduced aerobic glycolysis. *, p < 0.05, (n=10). (**C**) Principal component analysis (PCA) of bulk RNA sequencing data from eight AD donor-derived iSpheroid lines with and without Oxaliplatin treatment, demonstrating a distinct transcriptional response to treatment, (n=8). **(D)** Heatmap analysis showing the downregulation of glycolysis-associated genes following Oxaliplatin treatment. **(E)** Heatmap analysis demonstrating the upregulation of oxidative phosphorylation (OXPHOS)-related genes following Oxaliplatin treatment, indicating a shift toward enhanced mitochondrial respiration and oxidative metabolism. **(F)** Representative Seahorse XF Cell Mito Stress Test oxygen consumption rate (OCR) traces from control (CTRL) and oxaliplatin (OXA)-treated AD-derived iSpheroids. OCR was measured following sequential injections of oligomycin, FCCP, and rotenone/antimycin A, (n=2). **(G)** Quantification of basal respiration measured by Seahorse XF analysis in CTRL and OXA-treated AD-derived iSpheroids. *, p < 0.05. **(H)** Quantification of ATP-linked respiration measured by Seahorse XF analysis in CTRL and OXA-treated AD-derived iSpheroids. *, p < 0.05.

As elevated aerobic glycolysis, termed the Warburg effect, represents a hallmark of metabolic dysfunction in AD neurons^93^, we first quantified extracellular lactate production, as an indicator of glycolytic activity, following oxaliplatin treatment. Across iSpheroids generated from 10 independent AD donors, oxaliplatin treatment significantly reduced lactate production (**Figure 5B**; p < 0.05), indicating suppression of the Warburg-like metabolic phenotype previously observed in AD neurons^93^. These findings suggest that oxaliplatin shifts neuronal metabolism away from excessive aerobic glycolysis, supporting restoration of a more oxidative metabolic state.

To determine whether these metabolic changes were accompanied by an underlying transcriptional remodeling, we performed bulk RNA sequencing on iSpheroids from eight AD independent donors treated with oxaliplatin. Principal component analysis demonstrated clear separation between control and oxaliplatin-treated samples with about 83.7% variance explained in PC1 (**Figure 5C**), indicating that pharmacological treatment induced a robust and reproducible transcriptional response across genetically distinct AD donors. Rather than producing donor-specific effects, oxaliplatin consistently shifted the global transcriptomic landscape toward a common molecular state.

Because reduced lactate production suggested suppression of glycolytic metabolism, we next examined the expression of glycolysis-associated genes. Heatmap analysis demonstrated coordinated downregulation of glycolytic pathway genes following oxaliplatin treatment (**Figure 5D**), consistent with reduced reliance on aerobic glycolysis. In contrast, genes associated with oxidative phosphorylation (OXPHOS) were broadly upregulated (**Figure 5E**), indicating reciprocal activation of mitochondrial respiratory pathways. Together, these transcriptional changes demonstrated a coordinated metabolic transition from glycolysis toward oxidative metabolism rather than isolated regulation of individual metabolic enzymes.

To determine whether these transcriptional alterations translated into functional improvements in mitochondrial respiration, we performed Seahorse XF Cell Mito Stress Test analyses. Representative oxygen consumption rate (OCR) traces demonstrated increased respiratory activity following oxaliplatin treatment across the sequential injections of oligomycin, FCCP, and rotenone/antimycin A (**Figure 5F**). Quantification revealed a significant increase in basal respiration (**Figure 5G**; p < 0.05) and ATP-linked respiration (**Figure 5H**; p < 0.05) in oxaliplatin-treated iSpheroids, indicating enhanced mitochondrial function.

To further characterize the molecular response to treatment, we examined global transcriptional changes induced by oxaliplatin. Differentially expressed gene (DEG) analysis demonstrated widespread remodeling of the AD neuronal transcriptome following treatment, with numerous genes exhibiting distinct changes in expression (**Figure S17A**). These findings indicate that oxaliplatin does not simply regulate a small number of metabolic genes but instead induces broad transcriptional reprogramming consistent with restoration of neuronal metabolic homeostasis.

We next examined additional parameters of mitochondrial function derived from Seahorse analysis. Oxaliplatin treatment resulted in trends toward increased proton leak (**Figure S17B**), maximal respiration (**Figure S17C**), spare respiratory capacity (**Figure S17D**), and non-mitochondrial respiration (**Figure S17E**). Although none of these parameters reached statistical significance, all respiratory measurements showed a consistent directional increase after treatment, suggesting a global enhancement of mitochondrial bioenergetic capacity rather than selective modulation of a single respiratory parameter. The concordance between the metabolic analyses and transcriptomic data further supports the conclusion that oxaliplatin promotes metabolic remodeling toward improved mitochondrial function in AD iSpheroids.

Overall, these findings demonstrate that pharmacological intervention with oxaliplatin partially reverses key metabolic abnormalities associated with AD iSpheroids. Oxaliplatin reduced extracellular lactate production, suppressed glycolytic gene expression, induced oxidative phosphorylation pathways, and produced a consistent trend toward enhanced mitochondrial respiration across multiple functional parameters. Together with previous findings in 2D systems demonstrating an AD-associated Warburg effect^93^, these data establish this metabolic dysfunction as a reversible feature of AD neurons and iSpheroids as a scalable platform for evaluating therapeutic strategies, for instance, the ones aimed at restoring neuronal bioenergetic homeostasis.

### Identification of Trametinib as a pharmacological modulator of aging- and AD-associated phenotypes in human iSpheroids

Given the feasibility of the iSpehroids as a platform for therapeutical screening, we next used the system to identify compounds capable of modulating aging- and AD-associated neuronal phenotypes. Eleven candidate compounds under evaluation by the Longevity Consortium were independently applied to iSpheroids generated from three AD donors for two days, followed by bulk RNA sequencing to quantify drug-induced transcriptional responses (**Figure 6A-B**). AD donors were selected because their iSpheroids captured features of both advanced age and disease. This design enabled systematic comparison of compound-specific responses across independent donor backgrounds and provided a functional framework for identifying interventions that shift aged, diseased neurons toward a more favorable molecular state.

**Figure 6:**
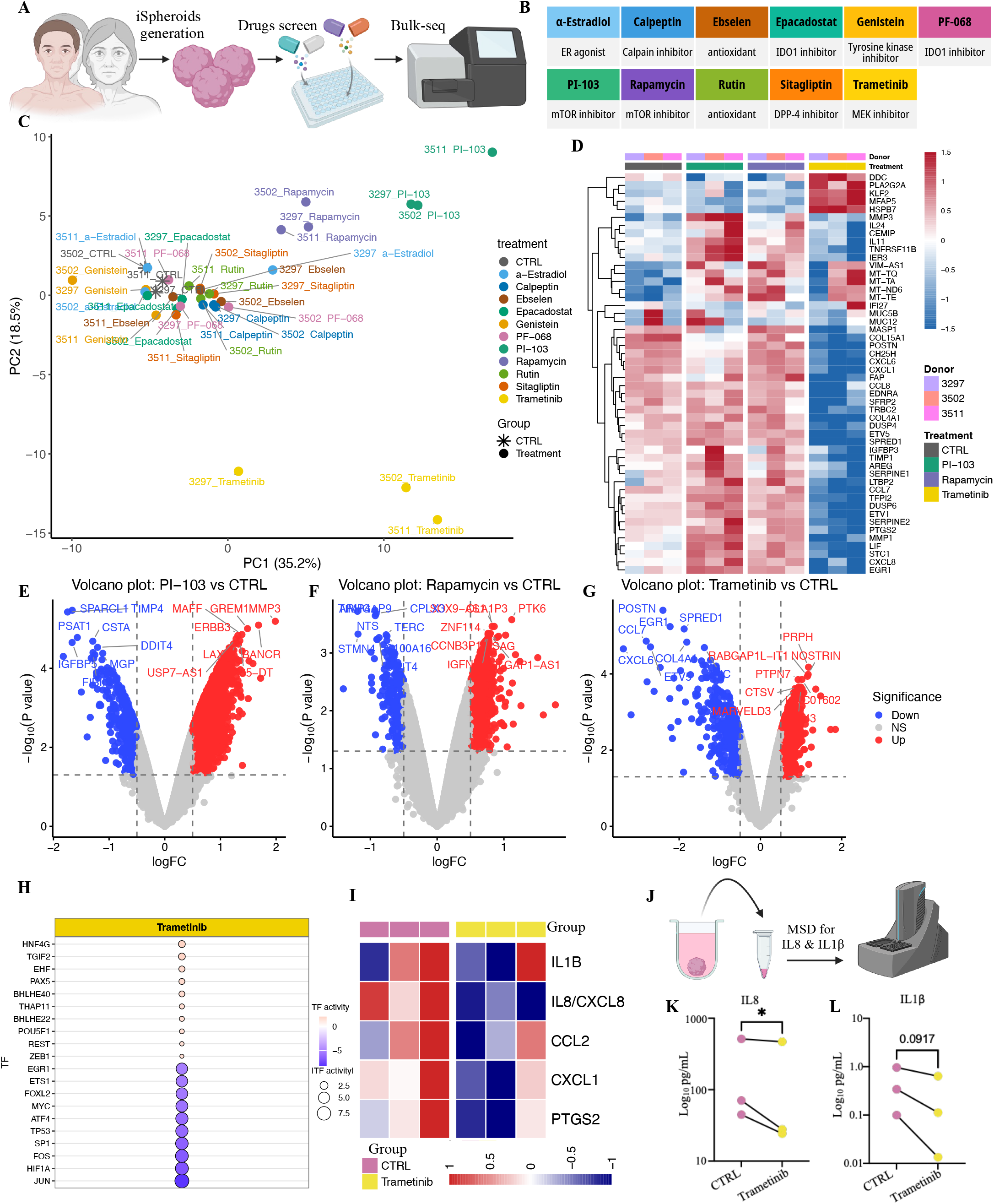
Identification and characterization of Trametinib as a modulator of aging- and AD-associated phenotypes in human iSpheroids. **(A-B)** Schematic overview of the drug screening workflow. Eleven candidate compounds were independently applied to iSpheroids derived from three AD donors for two days, followed by bulk RNA sequencing to assess drug-induced transcriptional responses. **(C)** Principal component analysis (PCA) of transcriptomic profiles demonstrating the effects of drug treatments across the three AD donor-derived iSpheroid lines, (n=3). **(D-G)** Heatmap of top 50 variable genes representing control samples and treated samples with PI-103, Rapamycin, and Trametinib (**D**), and volcano plots comparing control samples to PI-103, Rapamycin, and Trametinib (**E-G**). (H) Bubble plot of inferred transcription factor activities showing that Trametinib reduces the activity of inflammation-associated transcription factors while increasing the activity of transcriptional programs associated with pluripotency and neuronal function. **(I)** Heatmap demonstrating suppression of pro-inflammatory genes following Trametinib treatment. **(J-L)** Quantification of secreted pro-inflammatory cytokines, including IL-8 and IL-1β, measured by Meso Scale Discovery (MSD) assays (**J**), showing reduced cytokine secretion following Trametinib treatment. ns, p > 0.05 and *, p < 0.05 (**K-L**), (n=3).

Principal component analysis of the transcriptomic profiles showed that each compound induced a distinct molecular response across the three AD donor-derived iSpheroid lines (**Figure 6C**). Although several compounds produced relatively modest global changes, rapamycin, PI-103, and trametinib generated among the most pronounced transcriptional shifts and consistently separated treated samples from controls across donors (**Figure 6C-G**). Rapamycin and PI-103 inhibit mTOR signaling, a pathway extensively linked to longevity^129,130^. Trametinib, a MEK inhibitor, has likewise been associated with lifespan extension^131^, reduced inflammatory signaling^132^, and enhanced endogenous neurogenesis^133^. These observations motivated further analysis of trametinib as a candidate modulator of aging- and AD-associated neuronal states.

To define the regulatory programs underlying the trametinib response, we inferred transcription factor activity from the bulk RNA-seq data. Trametinib markedly reduced the activity of Generalized Stress Response (GSR)-associated transcription factors, including JUN, FOS, and ATF4, while increasing programs linked to neuronal identity and cellular rejuvenation, including PAX5 and OCT4 (**Figure 6H**). Consistent with these regulatory changes, trametinib also suppressed expression of proinflammatory genes including IL1B, IL8, CCL2, CXCL1, and PTGS2 (**Figure 6I).**

Because chronic GSR and neuroinflammation contributes to aging and AD progression^134–137^, we next asked whether suppression of inflammatory transcriptional programs translated into reduced cytokine secretion. Meso Scale Discovery (MSD) assays (**Figure 6J**) showed that trametinib significantly reduced secretion of IL-8 (**Figure 6K; p < 0.05**) and a trend toward a reduced IL-1β (**Figure 6L; p = 0.0917**) in conditioned medium from AD iSpheroids. Thus, the transcriptional response to trametinib was accompanied by a measurable reduction in extracellular proinflammatory signaling, supporting an anti-inflammatory effect at both molecular and functional levels.

To further define the molecular programs activated by trametinib, we performed STRING network analysis of pluripotency-associated genes upregulated after treatment. The resulting network was enriched for regulators of pluripotency, developmental plasticity, and cellular reprogramming, including POU5F1 (OCT4), NANOG, KLF4, PRDM14, ESRRB, LIN28A, LIN28B, DNMT3A, and DNMT3B (**Figure S18A**). MEK inhibition is known to support naive pluripotency^138^, and previous studies have shown that trametinib or related MEK inhibitors can activate OCT4 and other pluripotency-associated genes^139,140^. These findings raise the possibility that transient MEK inhibition increases cellular plasticity by engaging selected youthful or developmental transcriptional programs without complete dedifferentiation. We therefore asked whether trametinib also altered a donor-associated epigenetic age. Across 10 independent aged donors, methylation clock analysis showed a trend toward a modest reduction in iSpheroid PhenoAge that did not reach conventional statistical significance (**Figure S18B; p = 0.0819**). Although preliminary, this finding illustrates how the iSpheroid platform can connect drug-induced transcriptional remodeling with donor-specific aging readouts.

Finally, we compared inferred transcription factor activities across all compounds in the screen. Each compound generated a distinct regulatory signature, but trametinib uniquely combined suppression of inflammatory transcription factors with activation of neuronal differentiation and developmental regulatory programs (**Figure S18C**).

Together, these findings identify trametinib as a pharmacological modulator of aging- and AD-associated phenotypes in human iSpheroids. Trametinib remodeled transcription factor activity, suppressed GSR, pro-inflammatory gene expression, and pro-inflammatory cytokine secretion, and activated developmental and pluripotency-associated programs, accompanied by a modest reduction in measured PhenoAge. These results demonstrate that MEK inhibition can simultaneously perturb several measurable features of aged AD neurons and illustrate the utility of iSpheroids as a scalable platform for mechanism-oriented therapeutic screening.

## Discussion

The findings presented here establish iSpheroids as a scalable human neuronal platform that combines preservation of donor age with 3D structure, long-term neuronal maturation, disease phenotyping, and pharmacological perturbation. The platform also supports donor-specific glial integration and can be extended to non-human primate cells. Its defining feature is direct cellular conversion without passage through pluripotency, a process that can reset age-associated molecular and epigenetic signatures and thereby obscure donor age^50,61^. This distinction is particularly important for studies of aging and late-onset neurodegeneration, in which the biological history of the donor is an experimental variable rather than a feature to be erased. Across the analyses presented here, donor age and disease status remained detectable at multiple molecular and cellular levels, enabling interrogation of physiological aging and disease within a common experimental framework.

The reproducibility of neuronal generation across donors represents an important practical feature of the resource. Advanced donor age and AD did not compromise conversion efficiency, and iSpheroids remained predominantly neuronal across the 20 donors examined by snRNAseq. Moreover, iSpheroids supported long-term culture, electrophysiological activity, calcium dynamics, and synapse formation. These properties are important because preservation of aging alone is insufficient if the resulting cells cannot acquire or maintain the neuronal properties required to interrogate aging and late-onset neuronal phenotypes. The marked reduction in residual fibroblasts relative to conventional 2D direct conversion further suggests that the 3D environment influences neuronal conversion outcomes rather than simply providing an alternative geometry for culturing the same cells. This finding represents an important technical advantage, potentially reducing the need for FACS-based purification of converted neurons. One possible explanation for the improved neuronal conversion efficiency in iSpheroids is the extensive cell-cell contact established during spheroid compaction, which may promote contact inhibition and cell-cycle exit, thereby creating a cellular state more permissive to neuronal conversion^141^. In contrast, conventional 2D cultures may be less effective at maintaining such growth restraint, as ongoing cell loss creates unoccupied surface area whereas cell-plastic interactions permit residual fibroblasts to spread and proliferate. Thus, the physical organization of iSpheroids may actively favor neuronal conversion while restricting expansion of unconverted fibroblasts, contributing to the platform’s high neuronal purity.

The emergence of multiple neuronal states further indicates that iSpheroids are not transcriptionally homogeneous. Particularly notable was the identification of hippocampus-like populations expressing Prox1 and GRIK4, established markers associated with dentate gyrus^69,70^ and CA3^71^ identities, respectively. Given the vulnerability of hippocampal circuitry during aging and AD^65,142^, these populations create an opportunity to examine whether donor age or disease preferentially affects neuronal states relevant to vulnerable brain regions. At the same time, they should be interpreted as hippocampus-like transcriptional identities rather than complete representations of dentate gyrus or CA3 neurons. Establishing their relationship to bona fide hippocampal cell types will require deeper transcriptional comparisons, spatial characterization, and functional analysis. Differential patterning of iSpheroids using distinct media conditions may direct their differentiation toward specific cell types and/or brain regions.

Perhaps the most consequential property of iSpheroids for aging studies was demonstrated by methylation profiling. Whereas passage through pluripotency markedly reset epigenetic age^47,50,51,53^, directly converted iSpheroids retained age signatures resembling those of their parental fibroblasts. This finding provides molecular support for using direct conversion when donor age itself is central to the biological question. Importantly, however, preservation of age did not result in AD simply appearing as an exaggerated form of normal aging. Instead, AD iSpheroids exhibited an unexpectedly young-like epigenetic state relative to unimpaired aged donors. This observation is consistent with the broader concept that AD-associated epigenetic remodeling can diverge from normal aging rather than representing its linear acceleration. Thus, the distinction between young, unimpaired aged, and AD iSpheroids may reflect qualitatively different biological states rather than successive points along a single aging trajectory.

This distinction was reinforced across the transcriptomic and multi-omic datasets. Unimpaired aged iSpheroids did not consistently occupy an intermediate position between young and AD samples. Instead, they exhibited a distinct molecular state, particularly at the metabolic level. Lipidomic and metabolomic analyses showed comparatively lower intracellular lipid and metabolite abundance in unimpaired aged iSpheroids, whereas AD samples exhibited increased lipid abundance and metabolic remodeling. This finding is consistent with literature reporting that AD neurons accumulate lipids^113–115^. Proteomics similarly separated disease states and identified aging-associated proteins in unimpaired donors while revealing progressively stronger disease-associated changes in AD. Together, these observations raise the possibility that successful neuronal aging involves an actively maintained metabolic state that is distinct from both youth and AD. Whether the relatively low intracellular lipid and metabolite abundance observed in unimpaired aged neurons reflect metabolic efficiency, reduced biosynthetic demand, enhanced turnover, or another form of adaptation remains unresolved, but the convergence across independent omics measurements makes this an important direction for future investigation.

Importantly, the molecular differences observed in AD were accompanied by cellular phenotypes associated with neurodegeneration. Increased Fluoro-Jade C staining provided evidence of greater neuronal degeneration^89,90,92^, whereas changes in lactate were consistent with metabolic dysfunction^93^. Tau-associated abnormalities were apparent at multiple levels, including increased AT8 immunoreactivity, dystrophic bead-on-a-string neurites, and redistribution of phosphorylated Tau toward neuronal somata during prolonged culture. Detecting both 3R and 4R Tau further indicates that directly converted iSpheroids can support Tau isoform states associated with neuronal maturation. The progression of Tau-associated phenotypes with extended culture is particularly relevant because it suggests that long-term maintenance of donor-derived neurons can reveal disease features that may not be evident at earlier stages.

Aβ-associated phenotypes provided an additional pathological dimension. AD iSpheroids exhibited increased extracellular 82E1-positive deposits together with increased soluble Aβ42, whereas SEM independently revealed extracellular plaque-like structures. The convergence of antibody-based, biochemical, and ultrastructural measurements supports the presence of abnormal Aβ accumulation in AD iSpheroids. The appearance of Aβ deposits in AD iSpheroids is nonetheless notable because the AD lines examined here were not engineered to overexpress APP or familial AD mutations. This creates an opportunity to investigate mechanisms contributing to Aβ accumulation in naturally occurring donor backgrounds.

The metabolic abnormalities identified across these analyses also provided a disease phenotype that could be experimentally perturbed. Oxaliplatin reduced lactate production across multiple AD donors and induced a coordinated transcriptional shift characterized by reduced glycolytic gene expression and increased OXPHOS-associated programs. The direction of the Seahorse measurements was consistent with this transition toward increased mitochondrial respiration, although several individual respiratory parameters did not reach statistical significance. Accordingly, these findings are most appropriately interpreted as partial metabolic remodeling rather than complete restoration of neuronal metabolism.

The broader drug screen further demonstrated this capability. Distinct compounds generated separable transcriptional responses, with rapamycin, PI-103, and trametinib producing particularly pronounced changes across the AD donors examined. Trametinib was notable for simultaneously suppressing inflammatory transcriptional programs and increasing developmental and neuronal regulatory programs. Reduced JUN, FOS, and ATF4 activity was accompanied by decreased expression of inflammatory genes and reduced secretion of IL-8 and IL-1β, providing functional support for the transcriptionally inferred anti-inflammatory response. These results illustrate the value of combining transcriptomic screening with orthogonal phenotypic measurements rather than relying solely on differential gene expression to prioritize compounds.

The induction of pluripotency-associated regulators following trametinib treatment is particularly intriguing in the context of an age-preserving neuronal model. Transient activation of OCT4-, NANOG-, KLF4-, and related regulatory programs raises the possibility that aspects of cellular plasticity can be modulated after direct neuronal conversion without first returning cells to pluripotency. The accompanying reduction in PhenoAge across 10 aged donors was suggestive but did not reach conventional statistical significance. It therefore remains premature to interpret this response as neuronal rejuvenation. Instead, these observations identify a testable relationship between MEK signaling, developmental transcriptional programs, inflammation, and neuronal epigenetic age. Determining whether this state is transient or durable, whether neuronal identity remains stable after prolonged treatment, and whether molecular changes translate into improved neuronal function will be important next steps.

The ability to incorporate additional cell types further extends the experimental range of the resource. Integration of astrocytes and microglial lineage cells generated multicellular structures containing neuronal and glial populations, allowing the predominantly neuronal iSpheroid system to be expanded in a controlled manner. This modularity may be advantageous for distinguishing neuronal cell-autonomous phenotypes from effects that emerge through neuron-glia interactions. Similarly, survival and maintenance of neuronal identity following transplantation, together with host vascular infiltration, demonstrate compatibility with an in vivo environment. Extension of the conversion strategy to marmoset fibroblasts additionally suggests that the approach can support comparative studies across primate species. These applications are not intended to reproduce the complete cellular complexity of the human brain but instead provide experimentally tractable extensions of the same underlying platform.

Several limitations should be considered when interpreting these findings. iSpheroids are engineered neuronal structures and do not reproduce the full architecture, cellular diversity, vascularization, developmental history, or connectivity of the aged human brain. Although methylation analysis supports preservation of donor-associated age, aging is multidimensional, and future studies should determine how extensively additional chromatin, proteostatic, mitochondrial, and transcriptional features are retained. Likewise, the modest donor numbers used for several multi-omics and pharmacological analyses also constrain population-level inference. Finally, observations such as the young-like AD methylation state, the distinctive metabolic profile of unimpaired aged neurons, and the response of epigenetic age to trametinib should be viewed as hypotheses generated by the resources that require independent mechanistic validation rather than as established features of AD biology.

These limitations also define opportunities for subsequent use of the platform. Because the workflow is compatible with standardized multi-well culture and can be applied across genetically distinct donors, larger cohorts could be used to connect neuronal phenotypes with genotype, cognitive status, neuropathology, clinical progression, or therapeutic response. The same framework could be combined with spatial profiling, genetic perturbation, longitudinal imaging, additional glial populations, and higher-throughput pharmacological screening. In this setting, donor-to-donor variability becomes informative rather than simply experimental noise, potentially enabling identification of molecular features associated with resilience, vulnerability, or differential drug response.

Overall, the results support iSpheroids as a complementary resource positioned between conventional directly converted 2D neuronal cultures and pluripotent stem cell-derived 3D models, such as organoids. The platform does not aim to reproduce every feature of brain development or tissue organization; rather, its principal advantage is the ability to interrogate aged human neurons in a scalable 3D setting while retaining donor-associated biology. The distinctions observed between unimpaired and AD donors, the emergence of multiple AD-associated pathologies, and the ability to pharmacologically perturb measurable phenotypes collectively demonstrate the types of questions that can be addressed with this system. iSpheroids therefore provide a framework for studying how donor-specific neuronal aging relates to neurodegenerative vulnerability and for testing interventions directly in aged human neuronal backgrounds.

## Limitations of study

Although iSpheroids preserve donor-associated aging signatures and recapitulate multiple molecular and pathological features of sporadic AD, several limitations remain. Most AD phenotypes characterized here were examined in neuron-dominant iSpheroids and therefore do not capture the full contribution of astrocytes or microglia, or neuron-glia interactions to disease pathology. Although donor-specific astrocytes and microglia can be incorporated into iSpheroids, the effects of aging and AD on these populations were not investigated. Premixing astrocytes with fibroblasts also reduced neuronal conversion efficiency, suggesting that future studies should evaluate alternative integration strategies, including addition after neuronal conversion. More broadly, iSpheroids remain a reductionist in vitro model and do not reproduce the complete cellular, anatomical, vascular, or systemic complexity of the human brain. Neuronal conversion is driven by NGN2 and ASCL1, and the resulting populations, including neurons with hippocampus-like features, do not capture the full cellular diversity or spatial organization of their in vivo counterparts. Finally, the pharmacological experiments were designed to demonstrate phenotype-directed screening rather than to establish oxaliplatin or trametinib as candidate AD or aging therapies; their mechanisms, durability, and long-term effects will require further investigation. In addition, iSpheroids are not effective at capturing the very early characteristics and phenotypes of sporadic AD that iPSC derived Organoids strive to capture. On the other side, iSpheroids do capture characteristics of the specific individual from whom the fibroblasts were derived opening this type of *in vitro* model to a personalized phenotype.

## STAR★METHODS

### Key resource table

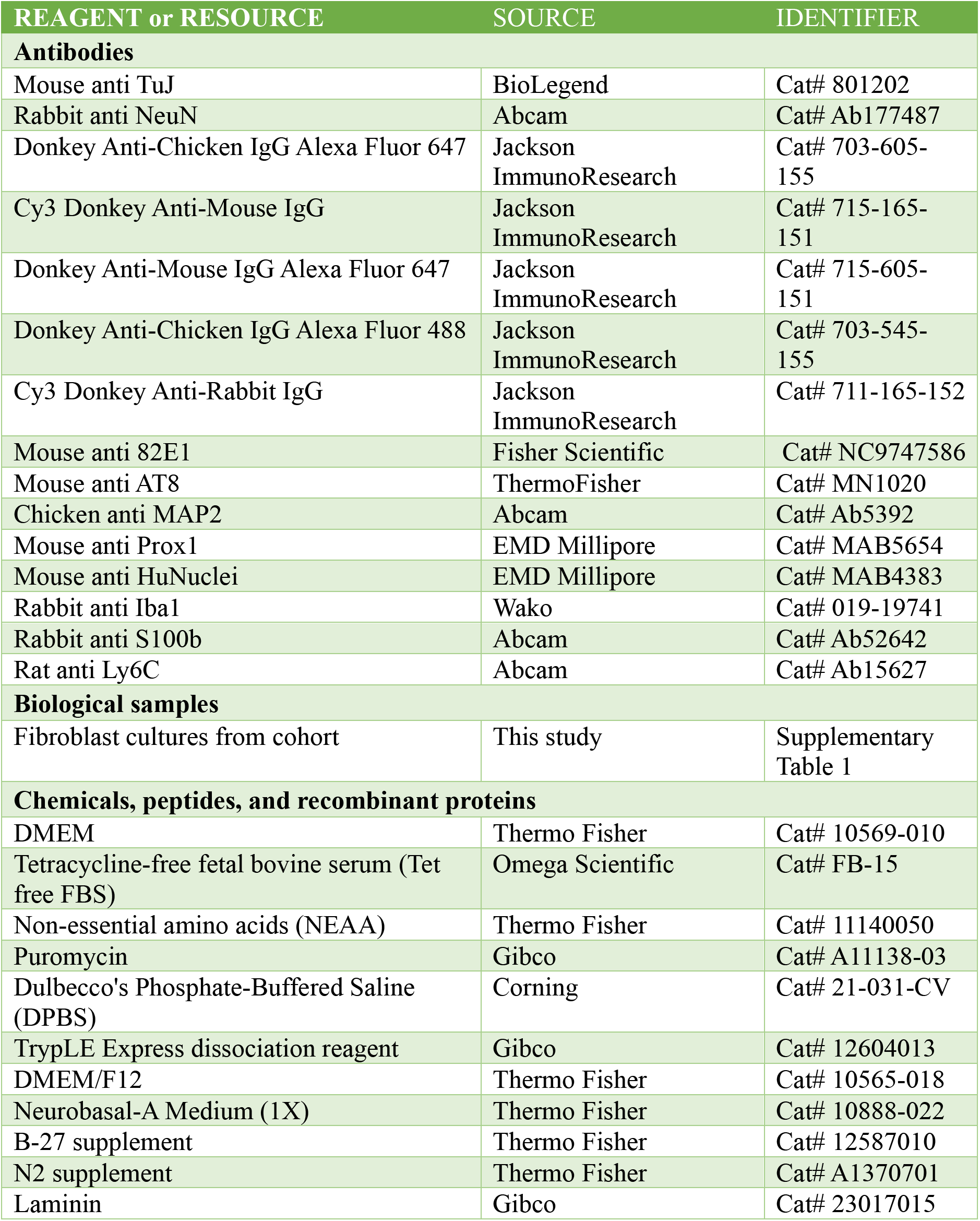
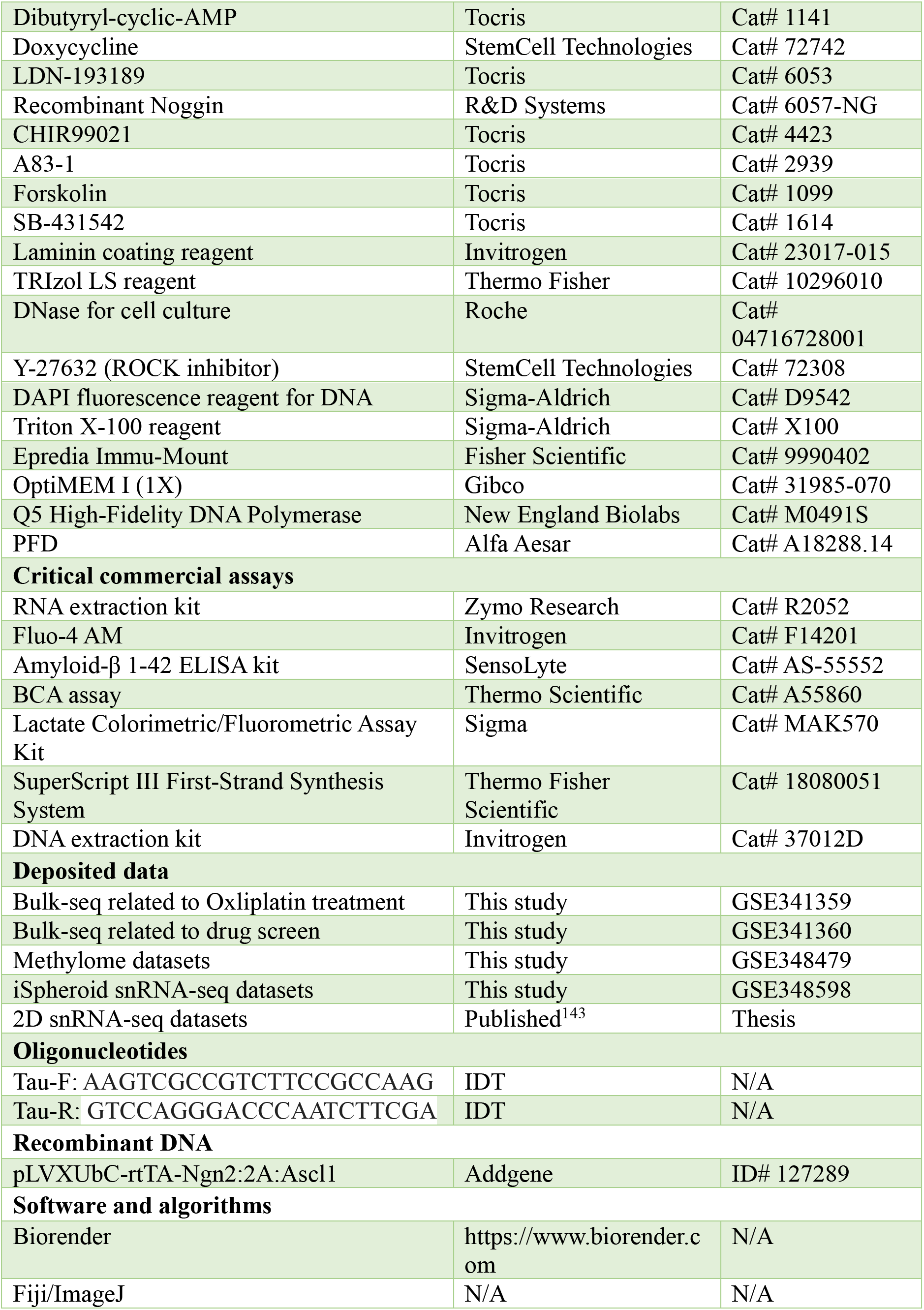
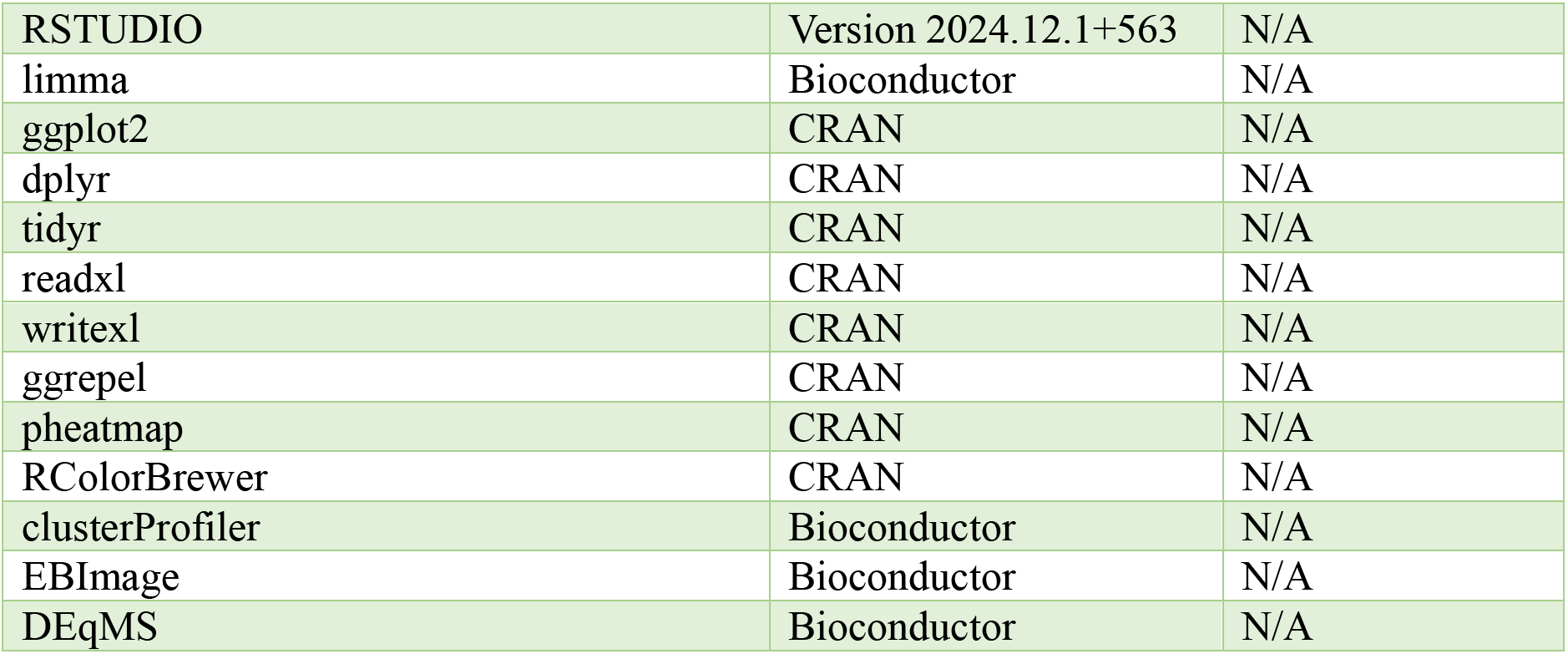

### Human cell culture

Human induced pluripotent stem cells (iPSCs) were maintained under feeder-free conditions with Matrigel (Cat# 354230) coating and mTeSR plus medium (Cat#100-0276). Primary human dermal fibroblasts were maintained at 37°C in a humidified incubator with 5% CO₂ in fibroblast low-serum medium consisting of Opti-MEM (Thermo Fisher Scientific, Cat# 31985088) supplemented with 2% TET-free fetal bovine serum (Omega Scientific, Cat# FB-01), 1× MEM non-essential amino acids (Thermo Fisher Scientific, Cat# 11140050), 20 ng/mL FGF2 (R&D Systems, Cat# 233-FB), and 1% penicillin/streptomycin (Thermo Fisher Scientific, Cat# 15070063). Cells were cultured on tissue culture-treated plates and passaged upon reaching approximately 80%–90% confluency. Cultures were washed once with phosphate-buffered saline (PBS, Cat# 14-190-144), dissociated using TrypLE Express (Cat# 12-604-013), collected by centrifugation, and resuspended in fresh fibroblast medium. Cell viability was determined by trypan blue exclusion dye using Countess Automated Cell Counters. Mycoplasma routinely performed in our laboratory and mycoplasma-negative cultures were used for subsequent experiments. All cell culture were performed in 37°C in a humidified incubator with 5% CO₂ with normoxia oxygen supply unless otherwise specified.

### Lentivirus production

HEK-293T cells were used to produce second-generation lentiviral particles. Briefly, HEK-293T cells were cultured to approximately 90% confluency and transfected with the lentiviral transfer plasmid together with second-generation lentiviral packaging plasmids using the Lipofectamine 3000 Transfection Kit according to the manufacturer’s instructions. Following transfection, lentivirus-containing supernatant was collected at two consecutive time points. The collected viral supernatants were purified through 0.45 um filters to remove cellular debris and concentrated by ultracentrifugation using a Beckman Coulter ultracentrifuge at 19400 RPM. Following centrifugation, the viral supernatant was carefully removed, and the concentrated lentiviral pellet was resuspended in an appropriate volume of basal culture medium and used for subsequent cell transduction or stored at −80 °C until use.

### Generation of fibroblast-derived iSpheroids

Human fibroblasts were maintained in fibroblast low-serum medium at 37°C and 5% CO₂. At confluency, fibroblasts stable for NGN2 and ASCL1 were dissociated using TrypLE Express, counted, and resuspended at a density of 1 × 10⁶ cells/mL. A total of 100 μL of cell suspension, corresponding to 1 × 10⁵ cells, was dispensed into each well of a 96-well round-bottom ultra-low-attachment plate using an eight-channel multichannel pipette. Cells were allowed to aggregate for 24 h to form spheroids, after which 150 μL of neuronal conversion medium was added to each well; this time point was designated day 0 of neuronal conversion. Neuronal conversion medium consisted of DMEM/F12 supplemented with B27 (10 mL/500 mL), N2 (5 mL/500 mL), GDNF (20 ng/mL), BDNF (20 ng/mL), cAMP (250 μg/mL), laminin (1 μg/mL), DMH-1 (2.5 μM), A83-1 (0.5 μM), CHIR99021 (3 μM), forskolin (5 μM), SB-431542 (10 μM), LDN-193189 (0.5 μM), ISX9 (20 μM), AlbuMax (0.5%), doxycycline (1 μg/mL), Antibiotic-Antimycotic (1%), and 1× MEM non-essential amino acids (MEM-NEAA). Beginning on day 3, cultures were fed every 3 days by aspirating 150 μL of medium at approximately a 45° angle using an eight-channel multi-well aspirator to minimize disturbance of the spheroids, followed by replacement with 150 μL of fresh neuronal conversion medium. On day 21, cultures were transitioned to neuronal maturation medium consisting of BrainPhys supplemented with B27 (10 mL/500 mL), N2 (5 mL/500 mL), GDNF (20 ng/mL), BDNF (20 ng/mL), cAMP (250 μg/mL), laminin (1 μg/mL), AlbuMax (0.5%), doxycycline (1 μg/mL), Antibiotic-Antimycotic (1%), and 1× MEM-NEAA, and were subsequently maintained with medium exchanges every 3 days. Wells located along the perimeter of the 96-well plates were excluded from experimental analyses because of increased evaporation and associated edge effects. Unless otherwise indicated, iSpheroids were used for downstream experiments beginning 60 days after initiation of neuronal conversion. The same procedure was used for the generation of marmoset fibroblast-derived iSpheroids.

### Astrocyte differentiation and iSpheroid integration

iPSCs from young, healthy aged, and AD donors were transduced with SOX9, NFIA, and NFIB (SNN) lentivector. Cells were cultured in astrocyte conversion medium consisting of DMEM/F12/GlutaMAX supplemented with 1% N2, 2% B27, SB431542 (100 µL per 500 mL), LDN-193189 (50 µL per 500 mL), and doxycycline (DOX; 200 µL per 500 mL). Cells were maintained in astrocyte conversion medium for 2–4 days, until conversion was achieved, and were subsequently transitioned to astrocyte maturation medium (AM) supplemented with DOX for maturation. For integration into iSpheroids, 30,000 iAstrocytes were mixed with 70,000 fibroblasts transduced with UNA. The mixed cell populations were allowed to aggregate and form spheroids, after which neuronal conversion was induced to generate astroglia–neuronal iSpheroids.

### Erythroid myeloid differentiation (EMP) and microglia-iSpheroid integration

EMP differentiation has been performed as previously established^144,145^ with slight modifications. Briefly, embryoid bodies (EBs) were generated in 96-well U-bottom plates by dissociating pluripotent stem cells using TrypLE and dispensing 100,000 cells per microwell in 150 μL mTeSR Plus medium supplemented with 10 μM ROCK inhibitor. EBs were allowed to form for 24 hours before initiating hematopoietic differentiation and EMP generation. Basal hematopoietic differentiation medium was supplemented with FGF2 (50 ng ml−1, Joint Protein Central), BMP4 (50 ng ml−1, Proteintech), Activin A (12.5 ng ml−1, R&D), ROCK inhibitor (1 μM, Stem Cell Technologies), and LiCl (2 mM, Sigma), and cultures were maintained under hypoxic conditions (5% CO2, 5% O2). Hematopoietic differentiation medium consisted of IMDM (50%, Thermo Fisher Scientific), DMEM/F12 (50%), ITSG-X (2% v/v, Thermo Fisher Scientific), L-ascorbic acid 2-phosphate magnesium (64 ug ml−1; Sigma), monothioglycerol (400 uM), PVA (10 ug ml−1; Sigma), GlutaMAX (1X, Thermo Fisher Scientific), chemically defined lipid concentrate (1X, Thermo Fisher Scientific), non-essential amino acids (NEAA, 1X, Thermo Fisher Scientific), and penicillin/streptomycin (P/S, 1% v/v, Thermo Fisher Scientific). On day 2, the medium was replaced and supplemented with FGF2 (50 ng ml−1, Joint Protein Central) and VEGF (50 ng ml−1, R&D), and cultures were returned to hypoxic conditions. On day 4, cells were transferred to normoxic conditions (5% CO2, 21% O2) and maintained in basal hematopoietic differentiation medium supplemented with FGF2 (50 ng ml−1, Joint Protein Central), VEGF (50 ng ml−1, R&D), TPO (50 ng ml−1, Proteintech), SCF (10 ng ml−1, Proteintech), IL-6 (50 ng ml−1, R&D), and IL-3 (10 ng ml−1, R&D) until day 10. At day 10, EBs were seeded onto Matrigel, where colonies were allowed to release erythroid myeloid progenitor cells (EMPs). EMPs were then co-cultured with iSpheroids at approximately 1,000 EMPs per iSpheroid in iSpheroid maturation medium supplemented with 25 ng ml−1 M-CSF (R&D), 100 ng ml−1 IL-34 (R&D), and 50 ng ml−1 TGFβ1 (R&D).

### Organoid differentiation

To generate embryoid bodies, intact PSC colonies were incubated with 1 mg ml−1 collagenase type IV (Invitrogen, 17104–019) for 20–45 min at 37°C and mechanically lifted using a cell scraper. Detached colonies were transferred to ultra-low-attachment dishes (Nunc) and maintained under stationary conditions for 2 days in mTeSR Plus supplemented with 10 uM ROCK inhibitor Y-27632 (Tocris Bioscience, 1254). Beginning on day 1, cultures were maintained in astrocyte medium (AstroM; ScienCell, 1800) containing astrocyte growth supplement (AGS; ScienCell, 1800), 2% FBS, 500 ng ml−1 Noggin (R&D Systems, 6057-NG), 10 ng ml−1 PDGF-AA (Proteintech, HZ-1215), and 1× 2-mercaptoethanol (Gibco, 21985032) through day 14.

On day 15, organoids were transferred to six-well ultra-low-attachment plates at 10–12 organoids per well and cultured on an orbital shaker at 75 rpm. Cultures were maintained for an additional week in AstroM supplemented with AGS, 10 ng ml−1 PDGF-AA, and 1× 2-mercaptoethanol. From day 22 onward, organoids were maintained at 75 rpm in DMEM/F-12 with GlutaMAX (Thermo Fisher Scientific, 10565018) supplemented with 1× N2 (Thermo Fisher Scientific, 17502–048), 1× B27 without retinoic acid (Thermo Fisher Scientific, 12587010), 1× MEM-NEAA (Thermo Fisher Scientific, 11140050), 10% FBS (Omega Scientific, FB-01; lot 991851), and 1× 2-mercaptoethanol.

For cultures maintained beyond day 60, the medium was additionally supplemented with 20 ng ml−1 BDNF (R&D Systems, 248-BDB), 20 ng ml−1 GDNF (R&D Systems, 212-GD), 0.2 mM ascorbic acid (STEMCELL Technologies, 72132), and 0.5 mM cAMP (Tocris Bioscience, 1141). Media were replaced every other day. Organoids at 3 months old were used for methylation clock analysis.

### Single-nucleus RNA library preparation and sequencing

Nuclei were isolated from freshly dissected iSpheroids according to the 10x Genomics protocol CG000124 (Rev E, Nuclei Isolation from Cell Suspensions and Tissues for scRNA-seq), with minor modifications. Briefly, freshly dissected iSpheroids were pooled and lysed in 1 ml of freshly prepared, ice-cold lysis buffer containing 10 mM Tris–HCl (pH 7.4), 10 mM NaCl, 3 mM MgCl₂, and 0.1% NP-40 in nuclease-free water using a Dounce homogenizer. Nuclei were collected by centrifugation at 500g for 8 min, and the supernatant was discarded. The nuclei pellet was resuspended in freshly prepared, ice-cold nuclei wash and resuspension buffer consisting of PBS, 1% BSA, and RNase inhibitor (0.2 U μl−1), followed by filtration through a 35-μm strainer (Corning). Nuclei were centrifuged at 500g for 5 min and resuspended in nuclei wash and resuspension buffer for a total of two washes. Following the second wash, nuclei were resuspended in ice-cold nuclei wash and resuspension buffer, and samples were pooled at a concentration of 800–1,200 nuclei per μl. Approximately 29,000 nuclei per channel, targeting an estimated recovery of 20,000 nuclei per channel, were loaded onto a Chromium Next GEM Chip G (10x Genomics) and processed using the Chromium Controller to generate single-nucleus gel beads in emulsion (GEMs). snRNA-seq libraries were prepared using the Chromium Next GEM Single Cell 3′ GEM Library & Gel Bead Kit v4 (10x Genomics) according to the manufacturer’s instructions. Libraries from different samples were pooled and sequenced to a target depth of 20,000 reads per nucleus on a NovaSeq X Plus (Illumina), using 28 bases for read 1, 90 bases for read 2, 10 bases for the i7 index, and 10 bases for the i5 index.

### Bulk RNA sequencing and analysis

For each RNA extraction, 8–16 iSpheroids were pooled, and total RNA was isolated using the Direct-zol RNA MicroPrep Kit (Cat# R2052) according to the manufacturer’s instructions. Purified RNA samples were submitted to Plasmidsaurus for outsourced bulk RNA sequencing. RNA quality assessment, library preparation, sequencing, demultiplexing, FASTQ file generation, and initial data analysis were performed by Plasmidsaurus according to their standard bulk RNA-seq workflow. Processed sequencing data were subsequently imported into RStudio, and further downstream analyses were performed locally.

### *In vivo* mouse transplantation

Xenotransplantation studies were performed as previously described^145^. Briefly, mice were anesthetized with 5% (induction) and 1-2% (maintenance) isoflurane in oxygen. Mice body temperature was maintained at 37°C and fixed into a stereotactic frame. Lidocaine (0.5%) was injected subdermally above the skull. The skin above the skull was cut with a small incision and a ∼3mm diameter craniotomy was performed by drilling into the skull. The underlying cerebral cortex (retrosplenial cortex) was aspirated with sterile blunt cannula (centered at approximately 4 - 4.5 mm posterior and 1.5 mm lateral to bregma). Sterile saline was used throughout the procedure to irrigate the lesion and Gelfoam was used to slow the bleeding. Three iSpheroids were placed into the cavity and covered with a 5mm coverslip to create a cranial window. Following completion of the surgery, carprofen (5mg/kg, ip) and buprenorphine (0.1mg/kg, sc) were administered for analgesic relief and to prevent inflammation. The mice were allowed to recover in a temperature-controlled cage and were returned to their home cages. Only mice were clearly identifiable iSpheroids grafts were subsequently analyzed.

### Electrophysiology

iSpheroids were visualized using a fixed stage upright microscope (BX51W1, Olympus) equipped with infrared differential interference contrast optics. Recordings were performed using a MultiClamp 700B amplifier interfaced to a Digidata 1500B (Molecular Devices) and data were acquired using pClamp11 software (Molecular Devices). All recordings were performed at room temperature (22-24°C) with flow rates of 1 ml/min in artificial cerebrospinal solution (ACSF) containing 124 mM NaCl, 26 mM NaHCO3, 10 mM glucose, 3 mM KCl, 2 mM CaCl2, 1.25 mM KH2PO4 and 1 mM MgCl2 equilibrated with 95%O2/5%CO2. Patch pipettes were pulled from borosilicate glass and had resistances of 5-8 MΩ when filled with internal pipette solution (130 mM K-gluconate, 1 mM EGTA, 5 mM Na-phosphocreatine, 2 mM Mg-ATP, 0.3 mM Na-GTP, 10mM HEPES; pH 7.3). All recordings were from cells with stable series resistance and input resistance. Excitatory postsynaptic currents (EPSCs) were recorded in voltage clamp mode with cells held at −70 mV. Peak sodium currents were recorded in voltage-clamp mode from a holding potential of −60 mV. A series of 400-ms voltage steps ranging from −100 to +140 mV in 10-mV increments was applied. For each cell, the maximal transient inward current evoked across the voltage-step series was identified and reported as the peak sodium current. Resting membrane potential (RMP) was measured in current-clamp mode in the absence of injected current (I=0) for 30 sec and the average current was plotted. Evoked action potentials (eAPs) were recorded in current-clamp mode with cells held at −65 mV using bias current. To accommodate the broad range of resting membrane potentials and excitability observed among iNs, the amplitude of current-injection steps was scaled to each cell’s bias current. The initial current step was set equal to the magnitude of the bias current required to hyperpolarize the cell from its resting membrane potential to −65 mV, and the step increment was set to one-tenth of this value. Current injections consisted of 40 sequential steps, beginning with 10 hyperpolarizing steps and followed by 30 depolarizing steps. Thus, for a cell requiring −20 pA of bias current to maintain −65 mV, current injections began at −20 pA and increased in 2-pA increments. Data analysis of eAPs was performed using NeuroExpress software (https://github.com/attilaszuc/NeuroExpress-patch-clamp-analysis) and clampfit. Current voltage response analysis was performed using Clampfit software (Molecular Devices).

### Multi-electrode array recordings

Neuronal activity was assessed using a multi-electrode array (MEA) MaxWell2 recording system. Mature iSpheroids were transferred onto MEA plates, excess media is removed and covered with perfluorodecalin (PFD). PFD presses on iSpheroids to enable efficient MEA recordings. Spontaneous extracellular activities were recorded according to MaxWell2 standard recommended procedures.

For each well, the firing rate of every recording electrode was calculated as the number of detected spikes divided by the recording duration. An electrode was classified as active if its firing rate was ≥ 0.1 Hz^146^. Network activity was summarized per well as the percentage of active electrodes (active electrodes divided by the total number of recording electrodes in that well) and the mean firing rate (MFR), averaged across active electrodes^146^. Each well corresponded to one donor (young, n = 6; unimpaired aged, n = 4; AD, n = 6), with empty and fibroblast-only wells recorded as controls.

Group differences between young, unimpaired, and AD iSpheroids were assessed for each metric using two-sided Mann–Whitney U tests^147^; no significant differences were detected (p > 0.05). Analyses were performed in Python (NumPy, SciPy, Matplotlib).

### Imaging and immunofluorescence

iSpheroids were imaged as live tissues, as post-fixation whole mounts, or following cryosectioning. Unless otherwise specified, iSpheroids were fixed in 4% paraformaldehyde (PFA) for 1–2 h at room temperature, followed by three washes with PBS and incubation in 30% sucrose at 4°C for long-term storage. For cryosectioning, iSpheroids were removed from the sucrose solution, embedded in OCT compound in a cryomold, and sectioned at a thickness of 20 μm. Approximately 30 sections could typically be obtained from a single iSpheroid. To preserve tissue integrity, iSpheroids were maintained in 30% sucrose and sectioned on demand. For immunofluorescence staining, cryosections were permeabilized and blocked in PBS containing 0.2% Triton X-100 and 6% horse serum for 30 min at room temperature. Appropriate primary antibodies were diluted in blocking solution and incubated with sections overnight. Sections were subsequently washed three times with blocking buffer, followed by incubation with the appropriate secondary antibodies for 2 h at room temperature. Sections were then thoroughly washed with PBS, counterstained with DAPI, and mounted for imaging. Whole-mount immunofluorescence staining was performed using a similar procedure. Images were acquired using multiple imaging platforms, including a slide scanner, Olympus FV5000, and Leica STELLARIS 5. Image processing, analysis, and quantification were primarily performed using ImageJ and locally in RStudio using the EBImage package.

### ELISA and colorimetric assays

Conditioned media or RIPA-based iSpheroid lysates were collected for ELISA and colorimetric analyses. Where appropriate, total protein concentration was determined before analysis using a BCA assay (Thermo Fisher Scientific, Cat. No. A55860) for normalization. Amyloid-beta 1-42 ELISA and extracellular lactate colorimetric assays were performed according to the manufacturers’ protocols. Absorbance or fluorescence signals were measured using a microplate reader, and analyte concentrations were calculated from standard curves generated in parallel with each experiment.

### Tau isoform assessment

For each RNA extraction, 8–16 iSpheroids were pooled, and total RNA was isolated using the Direct-zol RNA MicroPrep Kit (Zymo Research, Cat. No. R2062) according to the manufacturer’s instructions. RNA was reverse transcribed into complementary DNA (cDNA) using the iScript Reverse Transcription Supermix for RT-qPCR (Bio-Rad, Cat. No. 1708841). PCR was subsequently performed using primers designed to detect the 3R and 4R Tau isoforms. PCR products were resolved on a 1% agarose gel and imaged using a Bio-Rad imaging platform. Acquired bands were quantified by densitometric analysis using ImageJ. For each sample, 3R and 4R Tau band intensities were summed to represent total Tau, and the relative abundance of each isoform was calculated as a percentage of total Tau and subsequently plotted.

### DNA methylation clock assessment

Genomic DNA was extracted using Dynabeads magnetic beads according to the manufacturer instructions (Cat# 37012D). Purified DNA samples were submitted to TruDiagnostic (https://www.trudiagnostic.com/) for bisulfite conversion and hybridization to a custom Illumina Infinium MethylationEPIC v2.0 BeadChip. Raw intensity data (IDAT files) were processed by TruDiagnostic using their in-house R/Bioconductor pipeline, with minfi and the IlluminaHumanMethylationEPICv2anno.20a1.hg38 annotation. Samples with a mean detection p-value >0.05 were excluded. Beta values were normalized using single-sample Noob (ssNoob) background and dye-bias correction, and replicate EPICv2 probes were collapsed to a single value per CpG. Horvath pan-tissue DNAm age^84^ and DNAm PhenoAge^85^ were calculated from the normalized beta values using the methyAge function of the Enmix R package. CpGs not represented on the array were assigned reference beta values (TruDiagnostic reference medians, or ENmix reference values where these were unavailable) across all samples.

### Drug screening

Drug screening experiments were performed using mature iSpheroids at two months of culture in 96 multiwell plates. A panel of 11 compounds previously tested and recommended by members of the Longevity Consortium was selected for evaluation. Compounds were prepared as concentrated stock solutions and diluted into neuronal maturation medium immediately before use. iSpheroids were treated for 48 h with the following compounds at the indicated final concentrations: Epacadostat (IDO1 inhibitor, 5 μM), PF-06840003 (IDO1 inhibitor, 5 μM), Sitagliptin (DPP-4 inhibitor, 1 μM), Rutin (ADC cytotoxin chemical, 5 μM), Ebselen (HIV inhibitor, 1 μM), Genistein (EGFR inhibitor, 5 μM), Calpeptin (calpain inhibitor, 1 μM), PI-103 (PI3K/mTOR inhibitor, 500 nM), 17α-estradiol (estrogen/progesterone receptor inhibitor, 1 μM), Rapamycin (mTOR inhibitor, 50 nM), and Trametinib (MEK1/2 inhibitor, 50 nM). Epacadostat, PF-06840003, Sitagliptin, Rutin, Ebselen, Genistein, Calpeptin, PI-103, Rapamycin, and Trametinib were obtained from Selleck Chemicals, while 17α-estradiol was obtained from Sigma-Aldrich (Cat. No. E8750). Following 48 h of treatment, iSpheroids were collected for downstream analyses, primarily bulk RNA sequencing.

### Proteomics

Two-month-old iSpheroids representing four independents young, healthy aged, and AD donors were snap-frozen in liquid nitrogen at 24 iSpheroids per donor. Frozen iSpheroids were submitted to the UCSD Proteomics Core for label-free proteomic quantification. Precursor peak areas were log2-transformed, protein groups containing missing values were removed, and samples were median-centered. Differential protein abundance was assessed between young, unimpaired aged, and AD groups using DEqMS, with summed spectral counts for each protein group incorporated as the variance covariate. Pairwise comparisons were performed for young versus unimpaired and unimpaired versus AD samples. Proteins with *p* < 0.05 and |log2 fold change| > 1 were considered differentially abundant and were visualized using volcano plots. Gene Ontology biological-process enrichment analysis was performed separately on proteins increased in each direction of the pairwise comparisons, and enriched terms were ranked according to false-discovery-rate-adjusted significance and displayed as −log10(FDR). Sample-level relationships were examined by principal component analysis of normalized protein intensities. For heatmap visualization, log2 intensities from protein accessions mapping to the same gene symbol were averaged, followed by gene-wise standardization (row Z-score). Hierarchical clustering was performed using Euclidean distance and complete linkage.

### Metabolomics and lipidomics

Cells were initially washed once to remove extracellular components. For washing, cells at room temperature were resuspended in 400 µL of phosphate-buffered saline (PBS, pH 7.4) and subsequently centrifuged at 800 ×g-1 for 30 seconds at 4°C^148,149^. After the wash, the cell pellet was processed for metabolite extraction. The extraction was performed by resuspending the washed cell pellet in 1 mL solution of cold methyl tert-butyl ether (MTBE, Fisher Chemical; LC-MS grade) and methanol (Fisher Chemical; LC-MS grade) (3;1; v:v). The suspension was incubated on a thermomixer at 4°C with mixing at 1000 rpm for 45 minutes to facilitate efficient metabolite release, and following incubation was added 650 µL solution of water (Fisher Chemical; LC-MS grade) and methanol (3;1; v:v). The sample was centrifuged at 15,000 ×g-1 for 5 minutes at 4°C and separated into two phases. 400 µL supernatant from upper phase were carefully collected in glass tube as a-polar for lipidomics, and 600 µL supernatant from bottom phase were collected in 1.5 ml microcentrifuge tube as polar for metabolomics. The a-polar supernatant was solidified in liquid nitrogen, and subsequently dried overnight in a vacuum pressure concentrator (LABCONCO), while polar supernatant was dried at 25°C for overnight in speedvac concentrator (CentriVap Concentrator of LABCONCO) until complete removal of the solvent. The resulting polar-dried extract was reconstituted in 550 µL of 50 mM sodium phosphate buffer solution (PBS) (NaH2PO4/ Na2HPO4, Sigma Aldrich) with pH 7.4 in D2O (Sigma Aldrich) containing 0.5 mM trimethylsilyl propionate-d4 (TSP-d4, Sigma Aldrich), which served as an internal standard reference. Suspended solution was vortexed till complete dissolution and centrifuged at 15,000 ×g-1 for 10 minutes at 4°C. 500 µL transparent supernatant was transferred into 5 mm NMR tube (Wilmad) for NMR analysis. A-polar dried extract was reconstituted in 600 µL of CDCl3 (Sigma Aldrich) containing 0.03 % (v/v) trimethyl-silane (TMS) and centrifuged at 15,000 ×g-1 for 10 minutes at 4°C. 550 µL transparent supernatant was transferred into 5 mm NMR tube for NMR analysis. In the case of media for polar, 40 µL of media was added in 160 µL of 50 mM PBS (pH 7.4) in D2O containing 1.0 mM trimethylsilyl propionate (TSP) and vortexed it for one minute and transferred in 3 mm NMR tube for NMR analysis. A Bruker 800 MHz AVANACE NEO NMR spectrometer equipped with Bruker TCI (1H/2H/ 13C) cryogenic probe (BrukerBioSpin, Rheinstetten, Germany) was used to record all NMR spectra. 1D 1H NMR spectra of polar-neurons-cells from the zgesgp experiment were recorded at temperature of 298 K for 512 scans with a 5 second inter-scan delay and a 2 seconds per-scan acquisition time, spectral width 16 ppm with processed at data points 51280. The total acquisition time for each sample was 1 hour, 1 minute and 58 seconds. Topspin 4.5.0 was used for spectral processing following the acquisition. All spectra were zero-filled to 262144 points, and a 0.3 Hz line broadening was used. Similarly, polar-neurons-media were recorded for 128 scans with a 5 second inter-scan delay and a 2 seconds per-scan acquisition time, spectral width 16 ppm with processed data points 51280. The total acquisition time for each sample was 15 minutes and 35 seconds. Topspin 4.1.4 was used for spectral processing following the acquisition. All spectra were zero-filled to 131072 points, and a 0.3 Hz line broadening was used. Identification of metabolites and its quantification were analyzed at Chenomx NMR Suite 10.1 (Chenomx Inc., Alberta, Canada)3, and statistical analyses were performed using the web server Metaboanalyst 6.0 (https://www.metaboanalyst.ca,). 1D 1H NMR spectra of a-polar of neuron cells from the zg experiment were recorded at temperature of 298 K for 512 scans with a 7 second inter-scan delay and a 3 seconds per-scan acquisition time, spectral width 20 ppm with processed at data points 96152. All spectra were zero-filled to 262144 points, and a 0.3 Hz line broadening was used to quantify by the integral area of signals. 1D 1H NMR spectra with assigned identified metabolites were illustrated in Adobe Illustrator 2022.

### MSD inflammation panel

Secreted inflammatory proteins were quantified using the Meso Scale Discovery (MSD) V-PLEX Proinflammatory Panel 1 Human Kit, a 10-analyte multiplex electrochemiluminescence (ECL) immunoassay. Supernatant media containing secreted proteins was transferred to the panel and run according to standard manufacturer’s protocols. Data was acquired on the MSD Q60M instrument and analyzed using MSD Methodical Mind software. Protein concentrations were calculated from standard curves included on each plate and normalized for total protein using the Pierce BCA Protein Assay Kit.

### Seahorse metabolic assay

Cellular bioenergetics were assessed on the Seahorse extracellular flux analyzer (XF Flex analyzer) using the Seahorse XF 3D Mito Stress Test Kit. Individual iSpheroids were plated onto the Seahorse XF Flex 24-well 3D organoid microplate in Matrigel and incubated in standard media overnight. During the day of the assay, the plate was equilibrated in Seahorse XF DMEM Media containing 10mM glucose, 1mM sodium pyruvate, and 2mM L-glutamine in a low CO2 incubator for 45 minutes before taking measurements on the Seahorse XF Flex Analyzer. Oxygen consumption rate (OCR) and extracellular acidification rate (ECAR) were measured under basal conditions following sequential injection of mitochondrial inhibitors (oligomycin, FCCP, and rotenone/antimycin A). Bioenergetic parameters were analyzed using Seahorse Wave software.

### Genotyping and Imputation

Donors were genotyped on Illumina SNP arrays (GSA-24v3-0 or InfiniumCoreExome-24v1-4). IDAT files were converted to VCF using the bcftools gtc2vcf plugin (https://github.com/freeseek/gtc2vcf), and arrays genotyped on hg19 were lifted over to hg38 using GATK LiftoverVcf (https://gatk.broadinstitute.org/hc/en-us/articles/360037060932-LiftoverVcf-Picard-). Per-sample VCFs were merged with bcftools merge (https://samtools.github.io/bcftools/bcftools.html). Genotype imputation wasperformed via the Michigan Imputation Server (v2.0.9; https://imputationserver.sph.umich.edu) using the 1000 Genomes Phase 3 WGS (deep) reference panel (hg38, all populations). Haplotype phasing was performed with Eagle (https://alkesgroup.broadinstitute.org/Eagle/) prior to imputation with Minimac4 (v4.1.6; https://genome.sph.umich.edu/wiki/Minimac4). Concordance across samples was verified with bcftools gtcheck (https://samtools.github.io/bcftools/bcftools.html#gtcheck).

### Sample Demultiplexing

Cells were demultiplexed to individual donors using Popscle demuxlet (https://github.com/statgen/popscle). Prior to demultiplexing, the imputed VCF was subset to the donors present in each pool and sorted to match the chromosome order of the Cell Ranger BAM using popscle_helper_tools (https://github.com/aertslab/popscle_helper_tools). Variants were further filtered to retain only biallelic SNPs with no missing genotypes across all samples, removing uninformative positions (homozygous reference or homozygous alternate in all samples). The Cell Ranger BAM was filtered to retain only reads carrying valid cell barcodes overlapping retained SNP positions. Pileups were generated with popscle dsc-pileup, and donor assignment was performed with popscle demuxlet using the filtered BAM, filtered VCF, and Cell Ranger cell-barcode whitelist.

Cells were retained for downstream analysis if classified as singlets, with a log-likelihood difference between the best and second-best singlet assignment greater than one, at least 500 detected genes, and fewer than 5% of UMIs mapping to mitochondrial genes.

### snRNA-seq data preprocessing and cluster annotation

We used R (v.4.4.3) and Seurat (v.5)^150^ to process the snRNA-seq data and perform the downstream analysis. To remove poor-quality cells, cells were excluded from subsequent analysis if they expressed < 200 or > 5,000 genes, and cells with > 20% of mitochondrial gene content. In total, we retained 55,235 cells in our cohort. Integration was performed using Harmony (via R, version 1.2.3) on 30 PCs, correcting batch effects by donor and sequencing run. High resolution cell clusters were identified using the Seurat functions FindNeighbors and FindClusters with resolution = 0.02 for 2 clusters, which were then called “cell type” and classified the cells into fibroblast or iN, which was associated with the expression of known markers of fibroblasts and neurons, and resolution = 0.7 to identify 7 subtypes, then named Fibroblasts and iN clusters 1 through 6.

We performed differential expression analyses using the DESeq2 package (v.1.46)^151^ to identify markers for each cluster and characterize the clusters based on enriched functional pathways. Pseudobulk profiles were created for each donor by cluster using the sum of raw counts. Only donors with at least 30 cells per cluster were included in the pseudobulk downstream analysis.

Lowly expressed genes were excluded and only genes with more than 10 counts in at least 2 donors were considered. Differential expression analysis was performed using the Wald method as implemented in DESeq2. We included the sequencing run as a covariate during the analysis. Marker genes were then selected based on log2 Fold-change > 1 and adjusted p-value < 0.01. The top 100 genes were used to perform functional enrichment analysis through Metascape^152^.

Aging and Alzheimer’s disease signatures in the transcriptome of iSpheroids at single-cell level To evaluate if the iSpheroids can recapitulate some features of aging and Alzheimer’s disease using transcriptome data, we used pseudobulk profiles, as previously described, to perform differential expression analysis of the snRNA-seq data by diagnosis, i.e., AD, healthy aged and young donors, in each cluster through DESeq2. We ranked the genes based on the unadjusted p-value (log10 transformed) and converted the values to positive numbers if the log2 Fold-change > 0, and to negative numbers if the log2 Fold-change < 0.

### Scanning electron microscopy

iSpheroids from 12 donors, comprising four Young, four Healthy aged/unimpaired, and four AD donors, with four spheroids per donor, were processed for scanning electron microscopy (SEM) using a high-contrast heavy-metal staining protocol. Spheroids were initially fixed for 5 min at 35°C in 0.15 M cacodylate buffer (Ted Pella, Cat. 18851), pH 7.4, containing 2.5% glutaraldehyde (Ted Pella, Cat. 18426), 2% paraformaldehyde (Electron Microscopy Sciences [EMS], Cat. 19202), and 2 mM CaCl₂, followed by an additional 2 hr fixation on ice and five 3-min washes in ice-cold 0.15 M cacodylate buffer containing 2 mM CaCl₂. Samples were incubated for 1 hr on ice in freshly prepared reduced osmium solution containing 1.5% potassium ferrocyanide, 0.15 M cacodylate, 2 mM CaCl₂, and approximately 1.33% osmium tetroxide, followed by five 3-min washes in ddH₂O at room temperature. Samples were then incubated for 20 min at room temperature in freshly prepared, 0.22-µm-filtered 1% thiocarbohydrazide (TCH; VWR International, Cat. T1136-25G), washed five times in ddH₂O, and incubated in 2% aqueous osmium tetroxide (EMS, Cat. 19190) for 30 min at room temperature. Following five additional ddH₂O washes, samples were incubated overnight at approximately 4°C in 2% aqueous uranyl acetate (EMS, Cat. 22400), washed, and dehydrated through a graded ethanol series (20%, 50%, 70%, 90%, and 100%; 10 min each), followed by three additional 10-min incubations in 100% anhydrous ethanol. Samples were infiltrated with Spurr’s resin using 25% resin in ethanol for 1 hr 40 min, 50% resin overnight, and 75% resin for 2 hr, followed by three successive incubations in 100% Spurr’s resin for 1.5 hr, 2 hr, and 2 hr 15 min before embedding in fresh resin. Embedded samples were polymerized at 50°C overnight followed by 60°C for approximately 3 days. Polymerized samples were ultrathin sectioned at 70 nm using a DiATOME diamond knife on a Leica UC7 ultramicrotome and collected onto silicon chips using a serial-section SEM approach adapted from Horstmann et al.^153^ Silicon chips were mounted onto aluminum specimen stubs using double-sided carbon tape and imaged using a Zeiss Sigma VP scanning electron microscope equipped with a Gatan OnPoint backscattered electron detector.

## Quantification and statistical analysis

All quantified results are presented with error bars and standard deviations. Replicates are noted in each figure legend.

## Data and code availability

All accessions to data generated in manuscript can be found in Key Resource Table. For reviewers, Bulk-seq data can be accessed using these secure tokens axizgamqdjanfcb and ylopcacgrvolvwv. Methylome reviewers’ token is mhqxywqwrpetlcl. snRNAseq reviewers’ token is qhmzicqavlmnhmd.

## Acknowledgment

We thank the Longevity Consortium (award # U19 AG023122) for supporting this research. We acknowledge support from the Salk Institute Waitt Advanced Biophotonics Core Facility (RRID:SCR_014838) with funding from NIH-NCI CCSG P30 CA014195, NIH-NIA San Diego Nathan Shock Center P30 AG068635, the Waitt Foundation, Shared Instrumentation Grant S10 MH124757 and CIRM SRL Grant INFR6.2 15440, and The Henry L. Guenther Foundation, Salk Flow Cytometry Core (RRID:SCR_014839), Salk STEMCORE (RRID:SCR_014850), Salk Genomics Core (RRID:SCR_014842 and SCR_014846), and the UCSD Proteomics Core; and the UCSD SOM Microscopy Core (NS047101, OD030505, OD036455). We acknowledge KAUST sponsored Ibn Rushed fellowship support to Samhan Alsolami.

## Author contributions

**Conceptualization:** S.A., F.H.G.; **Methodology:** S.A., M.W., T.S.S, U.S., B.K., L.M., M.H., N.S C.B., D.B, J.R.H.; **Investigation:** S.A., M.W., U.S., B.K., L.M., M.H., C.B., D.B., E.M., S.C., R.L., J.R.H.; **Formal analysis:** S.A., T.S.S., N.S., K.O., M.M., M.C., E.M., S.C.; **Resources:** R.A., K.Y.; **Data curation:** S.A., T.S.S., K.O., M.C.; **Visualization:** S.A., T.S.S., K.O., C.B., D.B., M.M., J.R.H.; **Writing – original draft:** S.A., F.H.G.; **Writing – review & editing:** S.A., F.H.G., M.W., T.S.S., U.S., B.K., L.M., M.H., N.S., C.B., D.B., K.O., R.A., K.Y., M.M., M.C., E.M., S.C., R.L., J.R.H., J.R.J., J.M., J.M.T., L.J., G.J.M.; **Project administration:** S.A., F.H.G.; **Supervision:** J.M., J.M.T., L.J., G.J.M., F.H.G.

## Supplementary Materials

Supplemental figures and tables are placed in a separate document.

## Artificial intelligence use statement

ChatGPT 5.6 Sol was used to identify grammatical errors in the initial draft of this article following published guidelines^154^. The authors subsequently reviewed and independently revised the text and take full responsibility for the final content.

---

**Figure S1: Morphological characterization, long-term stability, and transcriptomic analysis of human iSpheroids.**

**(A)** Representative brightfield images of mature iSpheroids generated from human fibroblasts. Scale bar is 520 um.
**(B)** Time-lapse imaging snapshots illustrating the aggregation and self-assembly of fibroblasts into spheroids during the first 24 hours of culture. Yellow arrows at 5 hours indicate the process of cells compaction. Scale bar is 100 um.
**(C)** Quantification of TUJ1-positive neurons across iSpheroids derived from young, healthy aged, and AD donors.
**(D)** Schematic overview and corresponding immunofluorescence analysis of long-term cultured iSpheroids maintained for 270 days, including healthy aged (unimpaired) and AD donor-derived iSpheroids. Scale bar is 130 um.
**(E)** Quantification of induced neuronal (iN) and fibroblast cell populations across 20 donors based on single-nucleus RNA sequencing (snRNA-seq) analysis.

**(F-L)** Gene Ontology (GO) enrichment analyses of the seven major cellular clusters identified by snRNA-seq, highlighting the biological processes and molecular pathways associated with each cluster.

**Figure S2: Representative marker genes defining fibroblasts and induced neuronal subclusters.**

(**A**) UMAP visualization showing six induced neuronal (iN) subclusters and a distinct fibroblast population.

(**B-H**) Feature plots showing representative markers of the fibroblast population and each iN subcluster: fibroblast-associated markers ADAMTS6, GLIS3, and SPATS2L (**B**); iN cluster 1 markers ADAM19, NEURL2, and ITPKB (**C**); iN cluster 2 markers CCDC39, ENTPD1, and ZCCHC24 (**D**); iN cluster 3 markers PLCG2, EGFL7, and MT-ND4L (**E**); iN cluster 4 markers ELOVL2, CNR1, and DAZL (**F**); iN cluster 5 markers NRCAM, SYNDIG1, and KCTD16 (**G**); and iN cluster 6 markers WNT4, CACNA1D, and GPRIN3 (**H**). In the feature plots, darker purple indicates higher normalized gene expression.

**Figure S3: Identification and validation of hippocampal-like neuronal populations within human iSpheroids.**

**(A)** Heatmap showing the expression of canonical hippocampus marker genes enriched in Cluster 4, identifying this population as hippocampus-like neurons based on single-nucleus RNA sequencing analysis.
**(B)** Immunofluorescence validation demonstrating a subset of neurons expressing established dentate gyrus marker of Prox1, confirming the presence of hippocampus-like neuronal identities within iSpheroids. Yellow arrows indicate examples of cells positive for Prox1. Scale bar is 60 um. **(C-D)** Pseudobulk expression of **Prox1** (**C**) and **GRIK4** (**D**) following subdivision of induced neuron (iN) cluster 4 into three transcriptionally distinct subclusters (4_0, 4_1, and 4_2). ns, p > 0.05.
**(E)** UMAP visualization showing subdivision of the original iN cluster 4 into three subclusters (4_0, 4_1, and 4_2) while maintaining the remaining fibroblast and iN populations.

**Figure S4: Transplantation, vascularization, and neuronal maturation of human neuronal iSpheroids in the mouse brain.**

**(A)** Schematic overview of the experimental design in which human fibroblasts were directly converted into neuronal iSpheroids, matured *in vitro*, and stereotaxically transplanted into the mouse brain.
**(B)** Representative immunofluorescence images of the transplanted human neuronal iSpheroid labeled with the human-specific nuclear marker HuNu (yellow), blood vessels (green), and DAPI (blue), demonstrating vascularization of the transplanted graft by host-derived blood vessels. Scale bar is 100 um.
**(C)** Higher-magnification images of the transplanted graft shown in panel (B), demonstrating infiltration and close association of host blood vessels with the transplanted human neuronal iSpheroid. Scale bar is 100 um.
**(D)** Representative immunofluorescence images showing co-localization of the human-specific nuclear marker HuNu (green) with the neuronal marker MAP2 (red), confirming neuronal differentiation and survival of the transplanted human cells following engraftment.
**(E)** Representative immunofluorescence images showing co-localization of HuNu (green) with the mature neuronal marker NeuN (red), demonstrating maintenance of mature neuronal identity by the transplanted human iSpheroid following transplantation into the mouse brain.

**Figure S5: Generation of non-human primate neuronal iSpheroids by direct neuronal conversion.**

**(A)** Schematic overview of the experimental design in which non-human primate skin fibroblasts were transduced with a neuronal (UNA) lentiviral vector to induce direct neuronal conversion, followed by spheroid formation and long-term maturation to generate neuronal iSpheroids.
**(B)** Representative brightfield image of a mature non-human primate neuronal iSpheroid demonstrating uniform spheroid morphology following direct neuronal conversion. Scale bar is 210 um.
**(C)** Representative immunofluorescence images of the neuronal iSpheroid stained with DAPI (blue) and the pan-neuronal marker TUJ1 (green). The merged image demonstrates robust neuronal conversion and a predominantly neuronal composition throughout the 3D iSpheroid. Scale bar is 60 um.

**Figure S6: Electrophysiological characterization of human iSpheroids using patch-clamp recording and multielectrode array analysis.**

**(A)** Schematic overview of iSpheroid generation followed by functional assessment using whole-cell patch-clamp electrophysiology and MaxWell2 multielectrode array (MEA) recordings.
**(B)** Representative current-clamp recording demonstrating evoked action potentials in iSpheroid-derived neurons.
**(C)** Representative voltage-clamp recording showing spontaneous synaptic activity in iSpheroid-derived neurons.

**(D-K)** Quantitative patch-clamp analyses of multiple electrophysiological parameters across neurons derived from 8 cells representing young, 13 cells representing healthy aged, and 9 cells representing AD donors. ns, p > 0.05 and *, p < 0.05.

**(L-M)** Representative recordings and quantitative analyses of network activity measured using the MaxWell2 MEA platform on 6 young donors, 4 unimpaired aged donors, and 6 AD donors, demonstrating functional electrical activity and neuronal network formation within iSpheroids. ns, p > 0.05.

(**N**) MEA based network bursts analysis showing examples of burst found in an AD and an unimpaired donor. Red arrows indicate the location of the neuronal burst.
(**O**) Representative image of scanning electron microscopy of synapse-like structure in iSpheroids. Scale bar is 1 um.

**Figure S7: Generation and characterization of donor-specific glia-integrated iSpheroids.**

**(A)** Schematic overview of the generation of glia-integrated iSpheroids containing donor-specific astrocytes and microglia. Astrocytes are incorporated by combining approximately 30% astrocytes with 70% fibroblasts prior to spheroid formation, followed by standard neuronal conversion and maturation. Microglia are subsequently introduced through the addition of erythro-myeloid progenitors (EMPs) at a defined ratio and cultured in a modified medium supplemented with microglia-supporting factors for approximately 14 days to promote microglial differentiation and integration within the iSpheroids.

**(B-C)** Representative immunofluorescence images demonstrating successful integration of donor-specific microglia and astrocytes within iSpheroids on 9 donors, as identified by the microglial marker IBA1 and the astrocytic marker S100B and GFAP, respectively. Scale bar is 130 um.

**(D)** UMAP visualization of single-nucleus RNA sequencing data from glia-integrated iSpheroids derived from nine donors spanning young, healthy aged, and AD donors groups, revealing neuronal, fibroblast, astrocyte, oligodendrocyte progenitor cell (OPC), and microglia-related cellular subclusters.
(E) A pie chart showing the percentage of neuronal and glial subclusters.

**Figure S8. Distribution of cell populations across donors and cluster-specific marker expression in glia-integrated iSpheroids**

(**A**) Stacked bar plot showing the proportion of cells assigned to each transcriptionally defined cluster across individual donors, demonstrating reproducible representation of fibroblast-like, neuronal, glial, and immune-like populations while highlighting donor-to-donor variability in cluster abundance.
(**B**) Relative proportions of identified cell populations across donors (top) and heatmap showing cluster-enriched gene expression across fibroblast-like, early neuron-like, excitatory neuron-like, inhibitory neuron-like, OPC-like, glial-like, microglia progenitor-like, microglia-like, and undifferentiated cell populations (bottom). Genes are grouped according to the cell cluster in which they show preferential expression.

**Figure S9: iSpheroids capture age- and disease-related phenotypes.**

(**A**) UMAP representation of snRNA-seq data from young, healthy aged/unimpaired, and AD donors, showing the distribution of cells across donor groups.
(**B**) Heatmap showing the top 200 group-informative genes across young, unimpaired,AD donors.
(**C**) Average neuronal composition in snRNA-seq cluster 4-6 comparing unimpaired and AD donors. *, p < 0.05.

**Figure S10: snRNA-seq analysis reveals molecular signatures of aging and AD.**

A. KEGG pathway enrichment analysis comparing unimpaired and young donors
B. SEM images highlighting mitochondria (green) and ER (pink) morphologies in unimpaired and AD. Scale bar is 1000 nm.

**(C-E**) Bar graphs showing known AD-related genes significantly upregulated in AD compared with unimpaired donors using log_2_ (normalized gene expression + 1): CDK5 (**C**), PTPN1 **(D),** PDK1 **(E)**. *, p < 0.05 and **, p < 0.01.

**Figure S11: snRNA-seq captures neurodegenerative signatures in AD donors.**

(**A**) Heatmap showing increased expression of apoptosis-related genes in AD compared with unimpaired donors.
(**B**) Gene set enrichment analysis showing enrichment of multiple apoptosis-related pathways and biological processes in AD compared with unimpaired donors.
(**C**) Microplate reader-based measurement of extracellular lactate levels in independent 5 unimpaired and 7 AD donors. *, p < 0.05.
(**D**) Gel-based assessment of 3R and 4R Tau isoforms across young, unimpaired, and AD donors, 3 donors per category, with ImageJ-based quantification of 3R and 4R Tau bands and comparison of 4R Tau levels among young (Y), unimpaired (NC), and AD donors. ns, p > 0.05 and *, p < 0.05.

**Figure S12. Representative 1D ^1H NMR spectra of apolar lipid metabolites extracted from iSpheroids.**

**(A–C)** Representative 1D ^1H NMR spectra of apolar lipid extracts from young (**A**), AD (**B**), and unimpaired (**C**) iSpheroids. Major lipid functional group resonances are annotated throughout the spectra.

**Figure S13. Representative 1D ^1H NMR spectra of polar metabolites extracted from iSpheroids.**

**(A-C)** Representative 1D ^1H NMR spectra over the chemical shift range of 0.70–4.80 ppm from Young **(A)**, AD **(B)**, and Unimpaired **(C)** iSpheroids.

**(D-F)** Representative 1D ^1H NMR spectra over the chemical shift range of 4.81–8.79 ppm from young **(D)**, AD **(E)**, and unimpaired **(F)** iSpheroids. Major polar metabolite resonances are annotated throughout the spectra.

**Figure S14. Metabolomic profiling of extracellular metabolites from induced neuronal cultures.**

**(A)** Principal component analysis (PCA) of extracellular metabolomic profiles from young, unimpaired, AD, and control media samples. Each point represents an individual sample, with clustering indicating similarities in metabolite composition.
**(B)** Heatmap of differentially abundant extracellular metabolites across young, unimpaired, AD, and control media samples. Columns represent individual samples grouped by condition, and rows represent metabolites. Color intensity represents row-wise Z-score–normalized metabolite abundance, with red indicating higher abundance and blue indicating lower abundance.

**Figure S15. Representative 1D ^1H NMR spectra of polar metabolites in conditioned media from iSpheroids.**

**(A-D)** Representative 1D ^1H NMR spectra over the chemical shift range of 0.70–4.80 ppm from Control (**A**), young (**B**), AD (**C**), and unimpaired (**D**) conditioned media collected from iSpheroid cultures. **(E-H)** Representative 1D ^1H NMR spectra over the chemical shift range of 4.81–8.79 ppm from Control (**E**), young (**F**), AD (**G**), and unimpaired (**H**) conditioned media collected from iSpheroid cultures. Major polar metabolite resonances are annotated throughout the spectra.

**Figure S16. Differential protein expression and pathway enrichment analyses.**

**(A)** Volcano plots showing differentially expressed proteins between young and unimpaired donors.
**(B)** Pathway enrichment analyses of proteins differentially upregulated in either young or unimpaired.
**(C)** Volcano plots showing differentially expressed proteins between unimpaired and AD donors.
**(D)** Pathway enrichment analyses of proteins differentially upregulated in either unimpaired or AD.

**Figure S17. Oxaliplatin treatment alters gene expression and mitochondrial respiration in AD-derived iSpheroids.**

**(A)** Heatmap showing differentially expressed genes between control (CTRL) AD and oxaliplatin (OXA)-treated AD-derived iSpheroids. Columns represent individual samples, and rows represent differentially expressed genes. Color intensity represents row-wise Z-score–normalized gene expression, with red indicating higher expression and blue indicating lower expression.

**(B-E)** Seahorse XF analysis of mitochondrial respiration following oxaliplatin treatment, showing proton leak (**B**), maximal respiration (**C**), spare respiratory capacity (**D**), and non-mitochondrial respiration (**E**). Oxygen consumption rate (OCR) is presented in pmol/min. ns, p > 0.05.

**Figure S18: STRING interaction network and drug-induced transcriptional responses in iSpheroids.**

**(A)** STRING protein–protein interaction network of pluripotency-associated genes following trametinib treatment. Nodes represent pluripotency-associated genes/proteins, with node color indicating the direction of the trametinib-associated transcriptional response, and edges representing known or predicted functional interactions.
**(B)** Quantification of methyl-clock based neuronal phenoAge scores following treatment with control (Ctrl) or trametinib. Assessment spanned 10 donors. Each point represents an individual donor. ns, p > 0.05 and *, p < 0.05.
**(C)** Inferred transcription factor (TF) activity following treatment with the indicated compounds. Dot color represents the magnitude and direction of TF activity, whereas dot size corresponds to TF activity. Transcription factors are ranked according to their inferred activity within each treatment condition.

**Table S1: This table display some of young, healthy aged, and AD donors used in this study.**

